# Singlet oxygen leads to structural changes to chloroplasts during degradation in the *Arabidopsis thaliana plastid ferrochelatase two* mutant

**DOI:** 10.1101/2021.07.19.452378

**Authors:** Karen E. Fisher, Praveen Krishnamoorthy, Matthew S. Joens, Joanne Chory, James A. J. Fitzpatrick, Jesse D. Woodson

## Abstract

During photosynthesis, chloroplasts can produce large amounts of reactive oxygen species (ROS), particularly under stressful conditions. Along with other nutrients, chloroplasts also contain 80% of a leaf’s nitrogen supply. For these reasons, chloroplasts are prime targets for cellular degradation to protect cells from photo-oxidative damage and to redistribute nutrients to sink tissues. Multiple chloroplast degradation pathways have been described and are induced by photo-oxidative stress and nutrient starvation. However, the mechanisms by which damaged or senescing chloroplasts are identified, transported to the central vacuole, and ultimately degraded are not well characterized. Here, we investigated the subcellular structures involved with degrading chloroplasts induced by the ROS singlet oxygen (^1^O_2_) in the *Arabidopsis thaliana plastid ferrochelatase two* (*fc2*) mutant. Using a three-dimensional serial-block face electron microscopy analysis, we show up to 35% of degrading chloroplasts in *fc2* mutants protrude into the central vacuole. While the location of a chloroplast within a cell had no effect on the likelihood of its degradation, chloroplasts in spongy mesophyll cells were degraded at a higher rate than those in palisade mesophyll cells. To determine if degrading chloroplasts have unique structural characteristics allowing them to be distinguished from healthy chloroplasts, we analyzed *fc2* seedlings grown under different levels of photo-oxidative stress. A clear correlation was observed between chloroplast swelling, ^1^O_2_-signaling, and the state of degradation. Finally, plastoglobule enzymes involved in chloroplast disassembly were shown to be upregulated while plastoglobules increased their association with the thylakoid grana, implicating an interaction between ^1^O_2_-induced chloroplast degradation and senescence pathways.

## Introduction

During photosynthesis, plants naturally produce reactive oxygen species (ROS) in their chloroplasts (differentiated plastids that perform the photosynthetic reactions). Singlet oxygen (^1^O_2_) can be generated at photosystem I and superoxide and hydrogen peroxide can be generated at photosystem I (Asada 2006). Under mild conditions, these ROS can be quenched chemically (e.g., carotenoids and tocopherols quench ^1^O_2_) or enzymatically (e.g., chloroplast-localized 2-cysteine peroxiredoxins and thylakoid ascorbate peroxidase reduce hydrogen peroxide to water) (Foyer 2018). However, under various types of stress, including excess light (Triantaphylides et al. 2008), drought (Chan et al. 2016), salinity (Suo et al. 2017), and pathogen attack (Lu and Yao 2018), chloroplast ROS generation is increased and has the potential to overwhelm these protective mechanisms. Under such conditions, excess ROS can lead to the oxidation of DNA, proteins, and lipids, leading to photoinhibition and/or cell death (Triantaphylides et al. 2008). Therefore, chloroplasts must be in constant communication with the cell to mitigate ROS production and maintain efficient photosynthesis. This is achieved, in part, by the ROS themselves, which can act as both direct and indirect signals (Dogra and Kim 2019; Woodson 2019). Most work on such chloroplast signaling has focused on the numerous “retrograde signals” that broadly regulate nuclear gene expression in accordance with chloroplast development and stress (Chan et al. 2015; de Souza et al. 2017).

Recent work, however, has shown that some chloroplast signals can also induce chloroplast degradation (Woodson 2019). In such cases, chloroplast materials are sequestered into multiple types of structures for their delivery to lytic compartments such as the central vacuole (Zhuang and Jiang 2019). While the mechanisms behind chloroplast degradation are not well understood, such signals may allow for the removal of select damaged organelles from the cell in order to maintain a healthy population of chloroplasts (up to 120 in an *Arabidopsis* mesophyll cell (Pyke and Leech 1994)) in order to reduce ROS production and to increase photosynthetic efficiencies. Originally, chloroplast degradation was thought to be non-selective and utilized during senescence (Weaver and Amasino 2001). As chloroplasts contain ∼80% of the leaf nitrogen content, such degradation would allow for the redistribution of nitrogen and other essential nutrients (Makino and Osmond 1991). Early evidence for this kind of chloroplast degradation was reported in transmission electron micrograph (TEM) images of senescing pumpkin cotyledons (Harnischfeger 1973) and *Arabidopsis thaliana* mutants defective in plastid protein import machinery (Niwa et al. 2004).

More recent work has focused on understanding how chloroplast degradation is induced and the mechanisms involved in recognizing chloroplasts and removing them from the cell. These processes are more complex than originally assumed and multiple routes exist for degrading chloroplasts in plant cells (reviewed in (Nakamura and Izumi 2018; Zhuang and Jiang 2019)). Several routes involve the cellular autophagy machinery (i.e., cellular “self-eating”) and the core autophagy (*ATG*) genes involved in autophagosome formation (the canonical double membrane structure that delivers cargo to a lytic compartment such as the central vacuole). Entire chloroplasts can be transported from the cytoplasm to the central vacuole via chlorophagy, an autophagosome-dependent process that can be induced by UV-B or excess light stress (Izumi et al. 2017; Nakamura et al. 2018). At least under excess light stress, this process resembles ATG-dependent microautophagy, where the autophagosome partially surrounds the chloroplast as it is pushed into the central vacuole. In *atg* mutants, this pathway is non-functional. During carbon starvation in the dark, chloroplasts are also degraded by what may be a similar autophagy pathway (Wada et al. 2009). Several ATG-dependent piecemeal degradation pathways that disassemble chloroplasts and their components are also used by cells. In each case, these pathways involve distinct cytoplasmic structures that shuttle chloroplast components to the central vacuole. Rubisco-containing bodies (RCBs) transport Rubisco and other soluble proteins (Chiba et al. 2003), small starch granule-like structures (SSGLs) transport starch granules (Wang et al. 2013), and ATG8-INTERACTING PROTEIN1 (ATI1)-plastid bodies contain stromal, thylakoid, and envelope proteins (Michaeli et al. 2014).

Chloroplast degradation also occurs through ATG-independent pathways. The nuclear-encoded chloroplast vesiculation (CV) protein promotes chloroplast turnover when induced by drought or salinity stress, leading to the formation of CV-containing vesicles in the cytoplasm (Wang and Blumwald 2014). In rice, this pathway has been demonstrated to be important in balancing nitrogen supplies during stress-induced leaf senescence (Sade et al. 2018). Piecemeal degradation can occur through senescence-associated vacuoles (SAVs) that contain stromal protein, but lack thylakoids. Such structures remain in the cytoplasm during senescence and do not interact with the central vacuole (Martinez et al. 2008; Otegui et al. 2005).

In another pathway, chloroplast ^1^O_2_ has been shown to lead to the degradation of whole chloroplasts in mesophyll cells. Such a pathway was demonstrated in the Arabidopsis *plastid ferrochelatase two* (*fc2*) mutant, which conditionally accumulates large amounts of ^1^O_2_ in its chloroplasts (Woodson et al. 2015). Under diurnal light cycling conditions, the chloroplast-localized tetrapyrrole biosynthetic pathway is activated at dawn. In *fc2* mutants, however, a bottleneck effect leads to the rapid accumulation of the intermediate protoporphyrin IX (Proto). Like other unbound tetrapyrroles, Proto is an extremely photosensitive molecule that leads to the production of ^1^O_2_ in in the light. Thus, under diurnal light cycling conditions, *fc2* mutants experience a burst of ^1^O_2_, which results in the degradation of chloroplasts and eventually cell death. Under permissive 24h constant light conditions, plants avoid cell death. However, a subset of individual chloroplasts are still degraded, presumably due to low levels of ^1^O_2_. In some cases, these degrading chloroplasts appear to protrude into the central vacuole in an otherwise healthy cell, suggesting a selective chloroplast quality control pathway has been activated.

Genetic suppressors of the *fc2* cell death phenotype have been isolated and are called *ferrochelatase two suppressor* (*fts*) mutants (Woodson et al. 2015). Importantly, some suppressors do not reduce tetrapyrrole or ^1^O_2_ levels, indicating that such cellular degradation is a genetically controlled signal. Three loci involved have been shown to affect chloroplast gene expression, suggesting a chloroplast-encoded protein or RNA may be necessary to propagate the signal. Two of these (*PENTATRICOPEPTIDE-REPEAT CONTAINING PROTEIN 30* and *MITOCHONDRIAL TERMINATION FACTOR 9*), likely affect the post-transcriptional regulation of specific transcripts (Alamdari et al. 2020), while *CYTIDINE TRIPHOSPHATE SYNTHASE 2* indirectly regulates chloroplast gene expression by supplying dCTP for DNA synthesis (Alamdari et al. 2021). A fourth locus was shown to encode the cytoplasmic E3 ubiquitin ligase Plant U-Box 4 (PUB4), indicating a role for the cellular ubiquitination machinery involved in degradation (Woodson et al. 2015). Indeed, chloroplasts become ubiquitinated in *fc2* mutants, which may be a possible mechanism by which damaged chloroplasts are marked for turnover.

While the upstream events involved in initiating ^1^O_2_-induced chloroplast degradation are being elucidated, the degradation machinery itself has not been identified. However, there are several reasons to indicate that it is distinct from light stress-induced chlorophagy described above. First, TEM images have shown structural differences between chlorophagy and ^1^O_2_-dependent chloroplast degradation. While chlorophagy involves the translocation of intact-looking chloroplasts to the central vacuole, degrading chloroplasts in *fc2* appear to in an advanced state of degradation before interacting with the central vacuole. These chloroplasts lack chlorophyll and most of their internal membrane structures (Lemke et al. 2021; Woodson et al. 2015). ^1^O_2_-dependent chloroplast degradation can occur within three hours of ^1^O_2_ production (Woodson et al. 2015), while chlorophagy requires one or three days after excess light or UV-B stress, respectively (Izumi et al. 2017; Nakamura et al. 2018). Second, recent work has suggested these two pathways are genetically distinct. Chlorophagy can proceed without PUB4 (Kikuchi et al. 2020), while ^1^O_2_-dependent chloroplast degradation can proceed in the absence of the core autophagy machinery (Lemke et al. 2021).

Here, we decided to further investigate the structures involved in ^1^O_2_-induced chloroplast degradation in to order understand their unique characteristics. Furthermore, we wanted to assess the type of chloroplast associated with these structures, to determine if there are structural markers associated with degradation, and if ^1^O_2_ signaling is required. Our results demonstrate that such chloroplast and vacuole interactions are much more common than previously estimated. Chloroplast degradation itself is associated with several changes in chloroplast structure caused by ^1^O_2_ signals, including chloroplast swelling and dynamic changes in plastoglobule (PG) position. Furthermore, an increase in PG size and senescence-related transcripts suggests an overlap between ^1^O_2_-induced chloroplast degradation and senescence pathways.

## Results

### In *fc2* mutants, a proportion of degrading chloroplasts interact with structures in the central vacuole

When grown under short day diurnal light cycling conditions (e.g., 8 hours light/ 18h dark), *fc2* mutants experience wholesale chloroplast degradation and eventual cell death due to high levels of Proto and ^1^O_2_. Cell death does not occur under permissive 24h constant light conditions, but a portion of the chloroplasts are still targeted for degradation (Woodson et al. 2015). To explain this phenotype, we measured steady state tetrapyrroles in shoot tissue of three-day-old seedlings grown under 24h constant light. As shown in Table 1, *fc2-1* and *fc2-2* mutants accumulated excess Proto (but not other upstream or downstream intermediates) compared to wt and complementation lines. Compared to wt, *fc2-1* mutants also had a mild increase (22%) of bulk ^1^O_2_ in cotyledons (Fig. S1). Together, these results suggest that *fc2* mutants still experience a mild level of photo-oxidative stress even under permissive 24h constant light conditions.

**Table 1.** Steady state levels of tetrapyrrole intermediates in *fc2* mutants. Shown are mean levels (± SE) of tetrapyrroles (pmoles/g fresh weight) in shoot tissue of three-day-old seedlings grown under 24h constant light conditions (n ≥ 2 independent extractions).

| **Tetrapyrrole** | **wild type** | ***fc2-1*** | ***fc2-1***<br>***FC2::FC2-YFP*** | ***fc2-2*** | ***fc2-2***<br>***FC2::FC2-YFP*** |
| --- | --- | --- | --- | --- | --- |
| Uroporphyrinogen III | 14.4<br>± 0.5 | 14.5<br>± 0.8 | 14.0<br>± 0.3 | 15.6<br>± 1.5 | 10.6<br>± 1.1 |
| Coproporphyrinogen III | 5.6<br>± 0.3 | 4.6<br>± 0.6 | 6.2<br>± 0.2 | 6.3<br>± 0.4 | 3.4<br>± 0.1 |
| Protoporphyrin IX | 107.0<br>± 18.4 | 373.7<br>± 64.4 | 56.5<br>± 37.8 | 649.3<br>± 140.8 | 69.7<br>± 47.45 |
| Mg-Protoporphyrin IX | 11.3<br>± 2.1 | 12.6<br>± 2.3 | 4.4<br>± 0.3 | 10.14<br>± 1.1 | 2.1<br>± 0.4 |
| Mg-Protoporphyrin IX monomethyl ester | 27.6<br>± 4.7 | 36.32<br>± 5.1 | 16.2<br>± 4.7 | 29.26<br>± 1.7 | 5.0<br>± 0.7 |

In seedlings grown under 24h constant light, degrading chloroplasts can be recognized by compressed grana structures, irregular shapes, and abnormal vesicle-like structures. Occasionally, such degrading chloroplasts can be observed protruding or “blebbing” into the central vacuole. To characterize these structures and understand their role in chloroplast degradation, we further investigated these events using transmission electron microscopy (TEM) to image three-day-old *fc2-1* mutants grown under 24h constant light conditions (Figs. 1a and S2a). A close-up of the contact point between the vacuole and the degrading chloroplast reveals a close association between their two outer membranes; the tonoplast membrane and envelope, respectively (Fig. 1b). No additional membranes (e.g., double membrane autophagosomes) can be observed, which is in agreement with the core autophagy machinery being dispensable for such structures (Lemke et al. 2021). The image is remarkable in that the degradation appears to be targeted; other proximal cytoplasmic components such as mitochondria appear healthy and unaffected (Fig. 1c).

**Figure 1:**
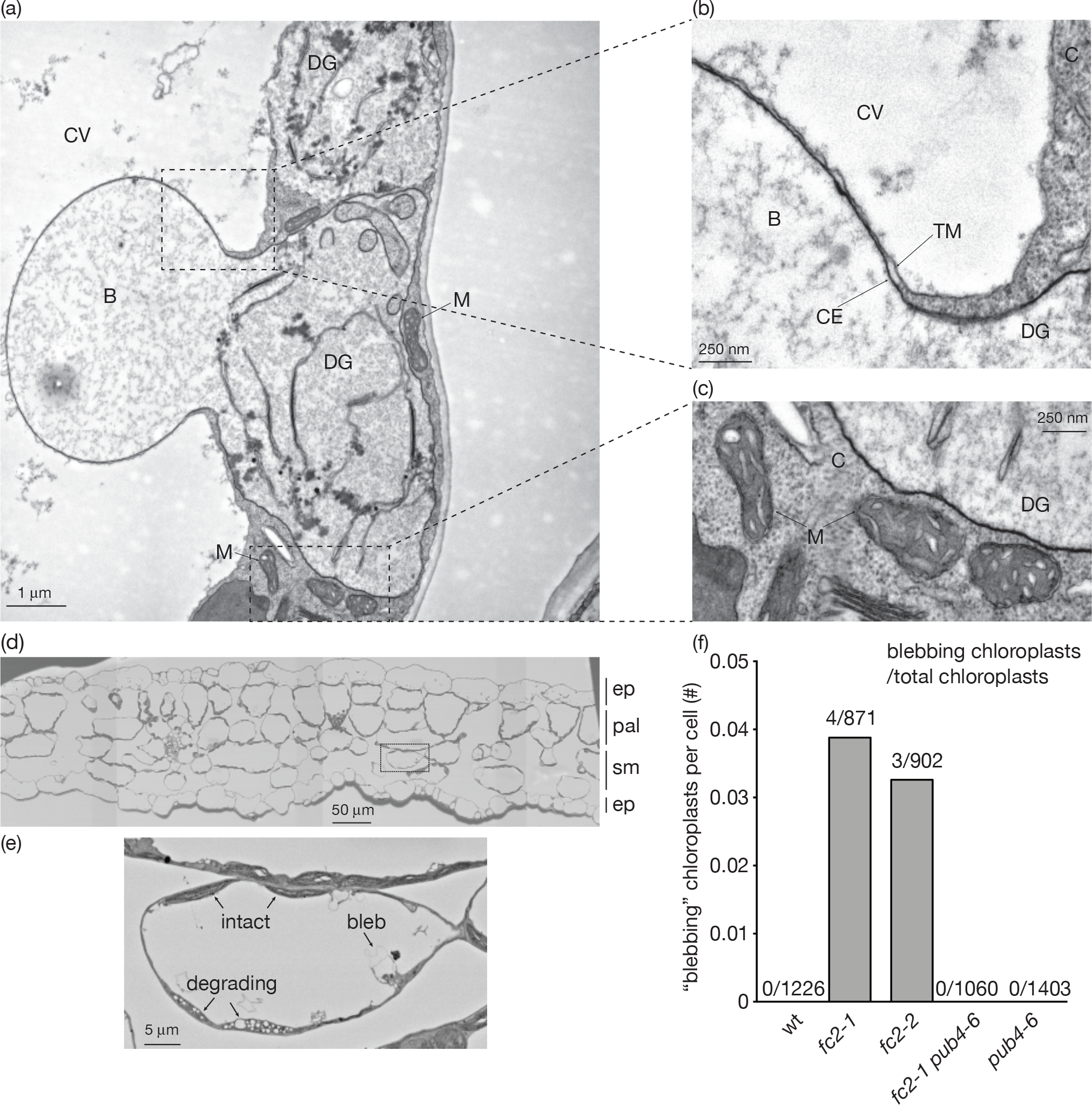
Quantification of chloroplast-vacuole interactions in *Arabidopsis fc2* mutants. Interactions between degrading chloroplasts and the central vacuole were analyzed using transmission electron microscopy (TEM). **A)** Shown is a representative two dimensional (2D) TEM image of a degrading chloroplast interacting with the central vacuole in a three-day-old *fc2-1* mutant grown under 24h constant light conditions. **B)** A zoomed-in image of the upper boxed area in panel A showing the interaction point between the tonoplast and degrading chloroplast envelope membranes. **C**) A zoomed-in image of the lower boxed area in panel A showing mitochondria in close proximity to the degrading chloroplast. Abbreviations: CV, central vacuole, DG, degrading chloroplast; M, mitochondria; B, chloroplast/vacuole bleb; CE, chloroplast envelope membrane; TM, tonoplast membrane. **D)** Shown is one FE-SEM tile-scanned cross section of a cotyledon from a five-day-old *fc2-1* seedling grown under 24h constant light conditions. Cell types are marked; ep, epidermal cells; pal, palisade cells; sm, spongy mesophyll cells. **E)** A zoomed in area marked on panel D showing intact, degrading and “blebbing” chloroplasts. **F)** Calculation of number (#) of blebbing chloroplasts per cell in the FE-SEM images (n ≥ 92 cells). Ratio above each bar indicates # blebbing chloroplasts/# total chloroplasts in images.

While degrading chloroplasts are commonly detected in two-dimensional (2D) TEM images of *fc2* mutants (on average, ∼19% of chloroplasts per cell) under 24h constant light conditions, blebbing structures are observed much less frequently (Lemke et al. 2021). To measure the rate of their occurrence, we returned to a previous FE-SEM tile-scanned data set of entire cotyledon cross-sections of five-day-old seedlings grown in 24h constant light conditions (Woodson et al. 2015). In these images, the epidermal, palisade mesophyll, and spongy mesophyll cell layers can be clearly identified (Fig. 1d). No single cell appeared to be experiencing cell death, and individual degrading and blebbing chloroplasts could be identified throughout the tissue sample (Fig. 1e). Using this data set, we previously reported rates of 0.15 (wt), 0.90 (*fc2-1*), 0.70 (*fc2-2*), 0.12 (*fc2-1 pub4-6*), and .03 (*pub4-6*) degrading chloroplasts per mesophyll cell (Woodson et al. 2015). Here, we surveyed blebbing chloroplasts in mesophyll cells and observed a similar pattern with much reduced frequency (Fig. 1f). Such structures were completely absent in wt, but the two independent *fc2* mutants (*fc2-1* and *fc2-2*) contained 0.04 and 0.03 blebbing chloroplasts per cell, respectively (or 0.46% and 0.33% of chloroplasts, respectively). In each case, the associated chloroplast also appeared to be degrading. No such structures were identified in the *fc2-1 pub4-6* mutant, consistent with PUB4 and ^1^O_2_ signaling being necessary for such chloroplast degradation (Woodson et al. 2015).

### A three-dimensional analysis of chloroplast-vacuole interactions

In order to obtain a spatial understanding of these structures and to determine the likelihood of those events amongst many chloroplasts contained within a single mesophyll cell, we looked to a three-dimensional (3D) Serial Block Face-Scanning Electron Microscopy (SBF-SEM) approach to provide a high-resolution scale covering a large 3D volume. Samples were prepared from five-day-old wild-type (wt) and *fc2-1* seedlings grown under 24h constant light conditions. Chloroplasts were then identified in cotyledon mesophyll cells and checked for degradation and any associations with vacuolar or blebbing structures. Compared to the 2D TEM images, a slightly larger percentage of degrading chloroplasts were observed in the SBF-SEM datasets. On average, 0.9% and 29.4% of chloroplasts were degraded in wt and *fc2-1* cells, respectively (Fig. 2a). In these SBF-SEM datasets, however, a much larger percentage of *fc2-1* chloroplasts were associating with “blebs” or structures that pushed into the central vacuole (∼8% of total chloroplasts (or ∼35% of degrading chloroplasts)). In wt cells, no such structures were observed out of 123 chloroplasts.

**Figure 2:**
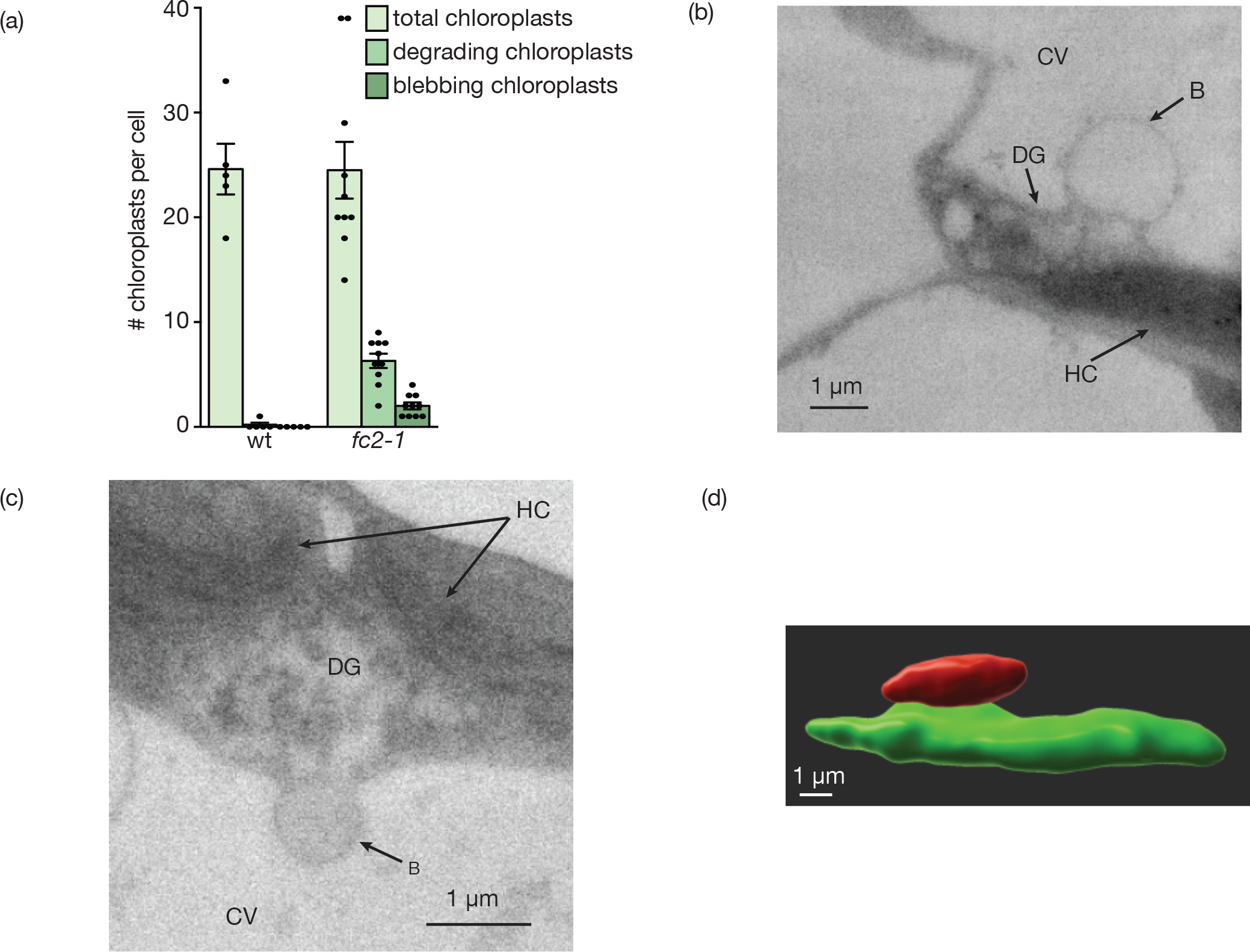
A three dimensional analysis of chloroplast degradation within the central vacuole. Chloroplast degradation in wt and the *fc2-1* mutant was analyzed using three dimensional (3D) Serial Block Face-Scanning Electron Microscopy (SBF-SEM). **A)** Shown are the mean number of total chloroplasts, degrading chloroplasts, and blebbing chloroplasts per cotyledon mesophyll cell observed using an SBF-SEM analysis of five-day-old seedlings grown under 24h constant light conditions (± SEM). In wt, 5 cells containing 123 chloroplasts were assessed. In the *fc2-1* mutant, 13 cells with 245 chloroplasts were assessed. Closed circles represent individual data points. To analyze the morphological characteristics of blebbing chloroplasts, representative structures were segmented and rendered out of the datasets. **B** and **C**) Shown are snapshots of two representative *fc2-1* chloroplasts used in the analysis. Abbreviations: CV, central vacuole, DG, degrading chloroplast; B, chloroplast/vacuole bleb; HC, healthy chloroplast. **D**) Shown is a 3D-rendering of the chloroplast depicted in panel B. Green indicates the cytoplasmic-localized chloroplast and red indicates the attached blebbing structure within the central vacuole.

To demonstrate the morphological characteristics of the degrading chloroplasts and associated structures, representative chloroplasts were segmented and rendered out of the datasets (Fig. 2b-d, Fig. S3a-d). The blebbing structures observed in *fc2-1*, were connected to the chloroplasts, but resided almost entirely within the vacuolar compartment. In some cases, proteinaceous matter could be detected near these contact points, suggesting a transfer of protein from the chloroplast to the central vacuole (Fig. 2c and Fig. S3d). Again, no such structures were observed associating with wt chloroplasts (Fig. S3b). The vacuolar structures had an average volume of 8.96 µm^3^ (13.88% of the chloroplast volume) and the average contact point between the chloroplast and the blebbing structure was 6.48 µm^2^ in area (Table 2). This small contact point may account for why the 2D TEM analysis underestimated the frequency of such interactions as these would often be missed while creating thin section samples for imaging. Finally, these calculations did not include one chloroplast-associated structure that was over an order of magnitude larger than the others (a volume of 177.6 µm^3^ and 202% of the associated chloroplast’s volume) (Fig. S3a, c, and d). It is unclear if this is a particularly large example of these chloroplast structures or a different type of structure altogether. Interestingly, this was the only blebbing chloroplast that was not in an obvious advanced stage of degradation.

**Table 2.**
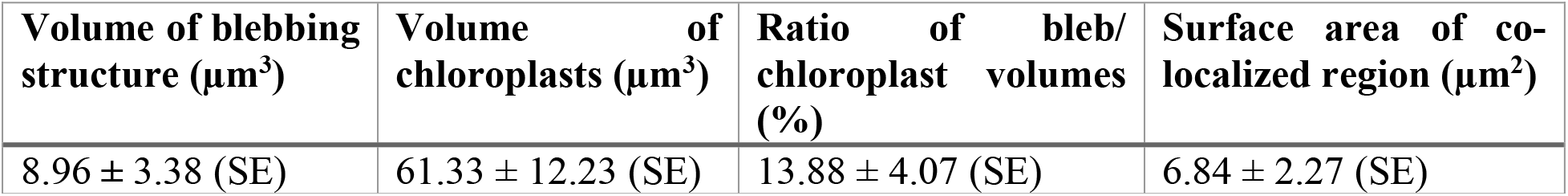
Measurements of degrading chloroplasts interacting with the central vacuole. Individual chloroplasts were segmented and rendered out of the three dimensional (3D) Serial Block Face-Scanning Electron Microscopy (SBF-SEM) data set of *fc2-1* seedlings. Shown are the average volumetric measurements of 10 chloroplasts associating with vacuolar blebbing structures. An eleventh chloroplast associating with an unusually large blebbing structure (177.6 µm^3^ and 202% of the chloroplast volume) was considered an outlier and not included in the analysis. Images of this chloroplast are in Fig. S3.

### Effect of cell type and chloroplast position on degradation

As described above, chloroplast degradation in the *fc2-1* mutant appears to occur in only a subset of chloroplasts when these seedlings are grown under 24h constant light conditions. Why only certain chloroplasts are targeted is unknown, but we hypothesized that the likelihood of a particular chloroplast being degraded may depend on cell type (palisade vs. spongy mesophyll) or position (adaxial vs. abaxial). To test this, we again returned to our *fc2-1* FE-SEM tile-scanned data. Individual degrading chloroplasts could readily be identified throughout the tissue sample by their compressed and disorganized thylakoid membranes (Fig. 1d). Chloroplasts were marked as intact or degrading in 75 mesophyll cells (both palisade (39 cells) and spongy (36 cells)) (Fig. 3a). We observed a slight increase in the rate of chloroplast degradation in spongy mesophyll cells, but this difference was not statistically different (p-value = 0.180, student’s t-test). However, it was noted that about half of cells had only intact chloroplasts (46% palisade and 61% spongy mesophyll cells). We performed this analysis again only using the cells with at least one degrading chloroplast (15 palisade and 20 spongy mesophyll cells) (Fig. 3a). In this case, there was a significant increase of chloroplast degradation in spongy mesophyll cells (p-value = 0.013, student’s t-test).

**Figure 3:**
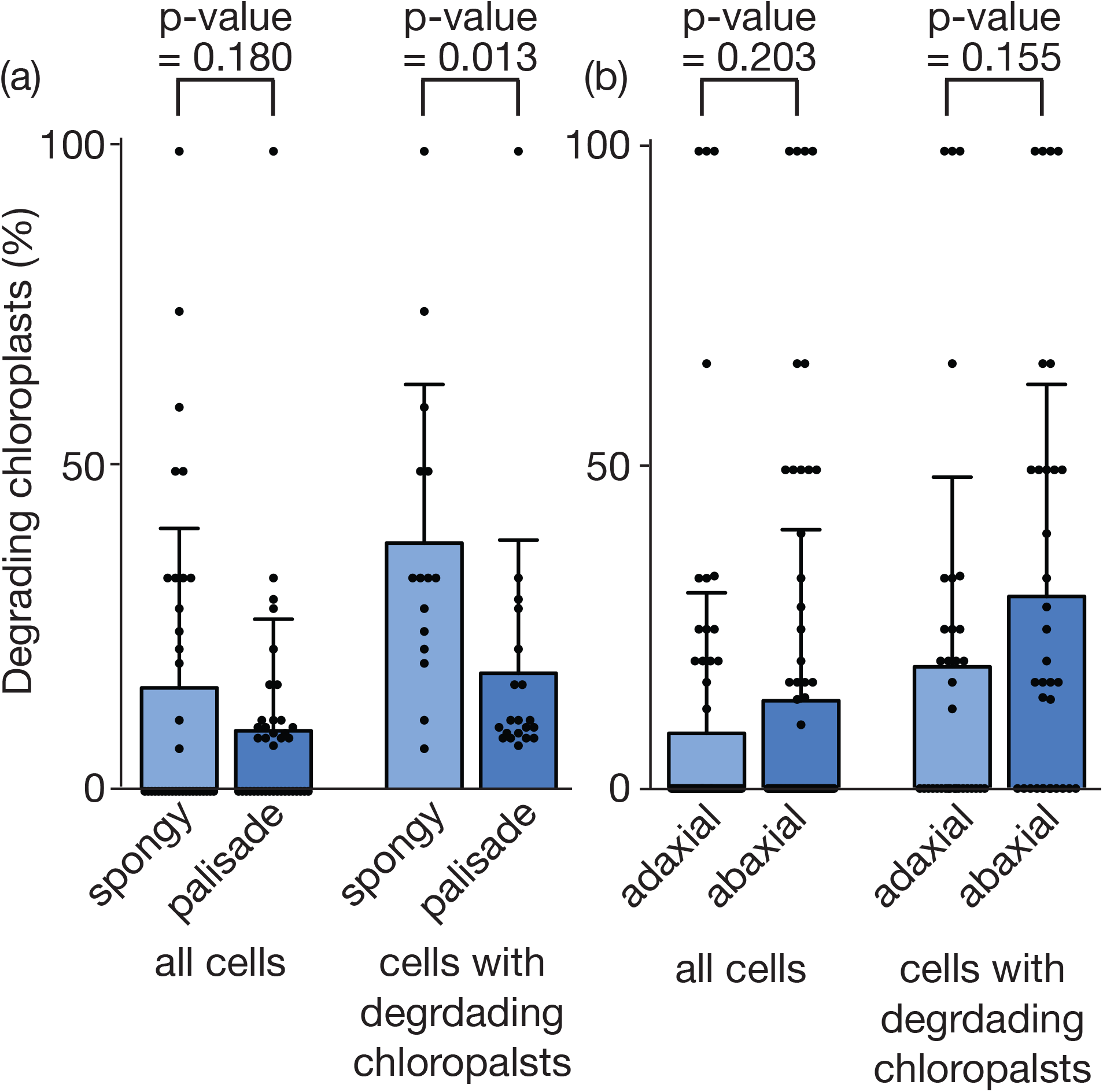
Effect of cell type and subcellular position on the frequency of chloroplast degradation. The frequency of chloroplast degradation in different cell types (palisade vs. spongy mesophyll) and subcellular positions (adaxial vs. abaxial) was assessed in FE-SEM images of five-day-old seedlings grown under 24h constant light conditions. **A)** Shown are the mean percentage (± SD) of degrading chloroplasts in the spongy mesophyll and palisade cells. **B)** Shown are the mean percentage (± SD) of degrading chloroplasts in the top (adaxial) versus bottom (abaxial) half of the cell. In both graphs, the left side of the graph includes all 75 cells (36 spongy mesophyll and/or 39 palisade cells), while the right side only includes the 35 cells (15 spongy mesophyll and/or 20 palisade cells) with at least one degrading chloroplast. Statistical analyses were performed by student’s t-tests. In all graphs, closed circles represent individual data points.

Next, we assessed if the position of a chloroplast within a cell, regardless of cell type, influenced its chances of being degraded. The same seventy-five cells analyzed above were divided in half at their equator and the number of degrading chloroplasts in both the top half (adaxial) and bottom half (abaxial) were counted. Again, we performed the analysis twice with all 75 cells or the 35 cells with at least one chloroplast being degraded (Fig. 3b). In both cases, we observed a slight increase in chloroplast degradation in the abaxial half of the cell, but it was not statistically significant in either case (p-values of 0.203 and 0.155 in all cells or only cells with degrading chloroplasts, respectively). Overall, the results suggest chloroplast degradation frequency may be partly influenced by cell type and is more common in spongy mesophyll cells. However, the position of the chloroplast within these cells does have a significant effect on their frequency of degradation.

### Chloroplasts in *fc2* mutants have reduced starch content

The above results suggest cells can target specific damaged chloroplasts for degradation in the *fc2-1* mutant. However, it is unknown how such organelles are recognized. One possibility is that ^1^O_2_-damaged *fc2-1* chloroplasts possess structural characteristics that allow them to be targeted. While chloroplast size has previously been shown to be reduced in *fc2-1* seedlings grown under 24h constant light (Alamdari et al. 2020), other ultrastructural characteristics have not been quantified. Therefore, to determine if there are morphological differences in these chloroplasts associated with degradation, we analyzed chloroplast ultrastructure by TEM in five-day-old seedlings of wt, *fc2-1*, *fc2-2*, *pub4-6* and *fc2 pub4-6* grown in 24h constant light (Fig. S2b).

We first assessed starch content in these mutants. Compared to wt, both *fc2-1* and *fc2-2* mutant chloroplasts contained significantly reduced starch compartment sizes (Fig. 4a). The majority of ellipsoidal chloroplasts contained at least one starch granule, suggesting that starch granule formation was not completely inhibited in the *fc2* mutants (Fig. 4b). Among the cells surveyed, all wt, *pub4-6*, and *fc2-1 pub4-6* chloroplasts contained at least one starch granule. In both the *fc2-1* and *fc2-2* mutants, however, several chloroplasts lacked starch granules (7% of *fc2-1* and 16% of *fc2-2*). This supports previous observations of delayed plant growth and development in *fc2* mutants (Scharfenberg et al. 2015; Woodson et al. 2015). Consistent with its role in suppressing *fc2* phenotypes, the *pub4-6* mutation was also able to partially restore starch content on the *fc2-1* mutant (Fig. 4a and b). This effect was specific to the *fc2-1* background, as the single *pub4-6* mutant had starch content equivalent to the wt control.

**Figure 4.**
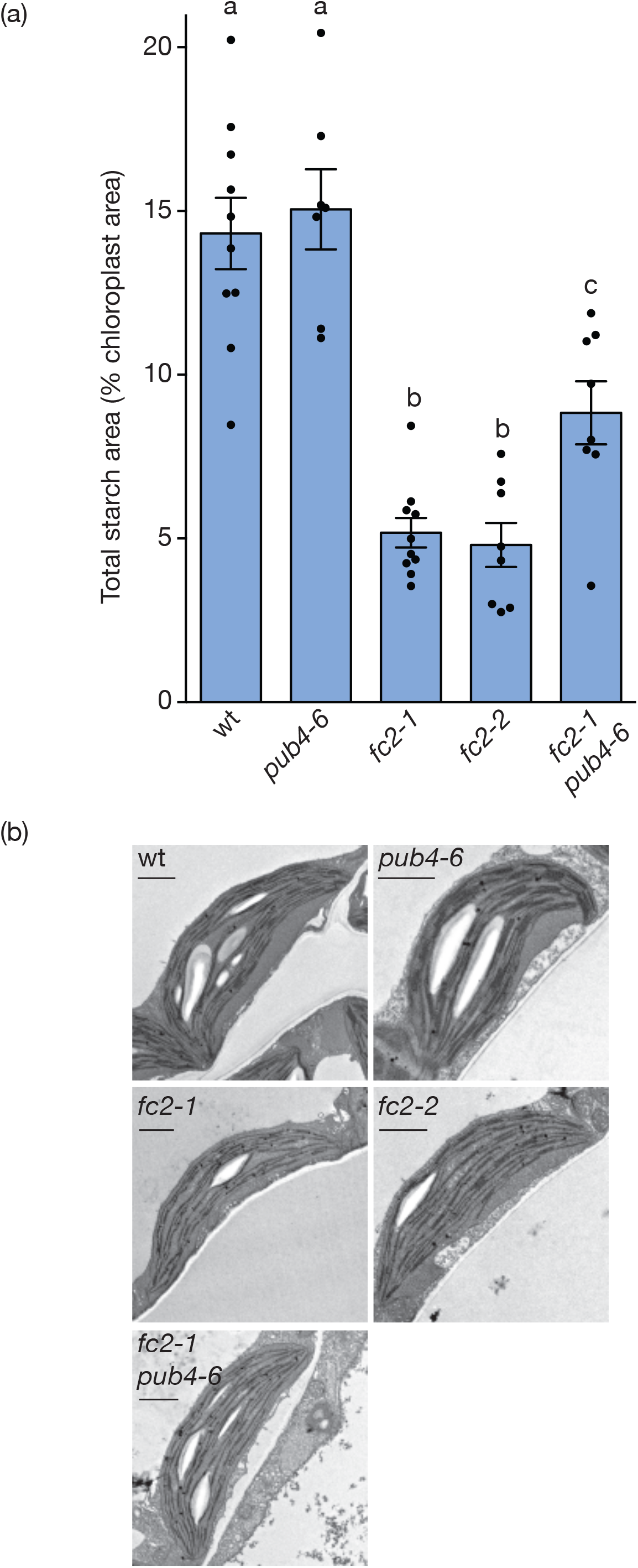
*fc2* mutants have reduced starch content. Starch content in the chloroplasts of *fc2* mutants was assessed using TEM. **A)** Shown is the mean starch granule area (± SEM) per chloroplast in five-day-old seedlings grown in 24h constant light (n ≥ 7 cells). Statistical analyses were performed by a one-way ANOVA test followed by a Tukey-Kramer post-test. Different letters above bar graph represent significant differences (p-value ≤ 0.05). Closed circles represent individual data points. **B)** Representative images of chloroplasts and their starch granule(s). Scale bar = 1µm.

### An increase in chloroplast swelling is associated with degradation

Swelling of chloroplasts and mitochondria has previously been suggested as a marker for stress that precedes chlorophagy (Nakamura et al. 2018) and mitophagy (Kim et al. 2007), respectively. Therefore, we wanted to examine if such swelling is also associated with ^1^O_2_-induced chloroplast degradation in *fc2* mutants under constant and cycling light conditions. To this end, we grew wt, *fc2-1*, *and fc2-1 pub4-6* seedlings under different 24h photoperiods (4h light/20h dark, 8h light/16h dark, 16h light/8h dark, and 24h light/0h dark), which has been shown to induce varying levels of photo-oxidative stress (Woodson et al. 2015). To assess the amount of cellular degradation induced by these treatments, we tested for cell death by staining seven-day-old seedlings with trypan blue. As expected, wt experienced no cell death under any conditions, while *fc2-1* seedlings exhibited significant cell death only under the two shortest photoperiods (4h light/20h dark and 8h light/16h dark) (Figs. 5a and S4). Under all conditions, the *fc2-1 pub4-6* mutant experienced no significant cell death, confirming that such cellular degradation was ^1^O_2_ signaling dependent.

**Figure 5.**
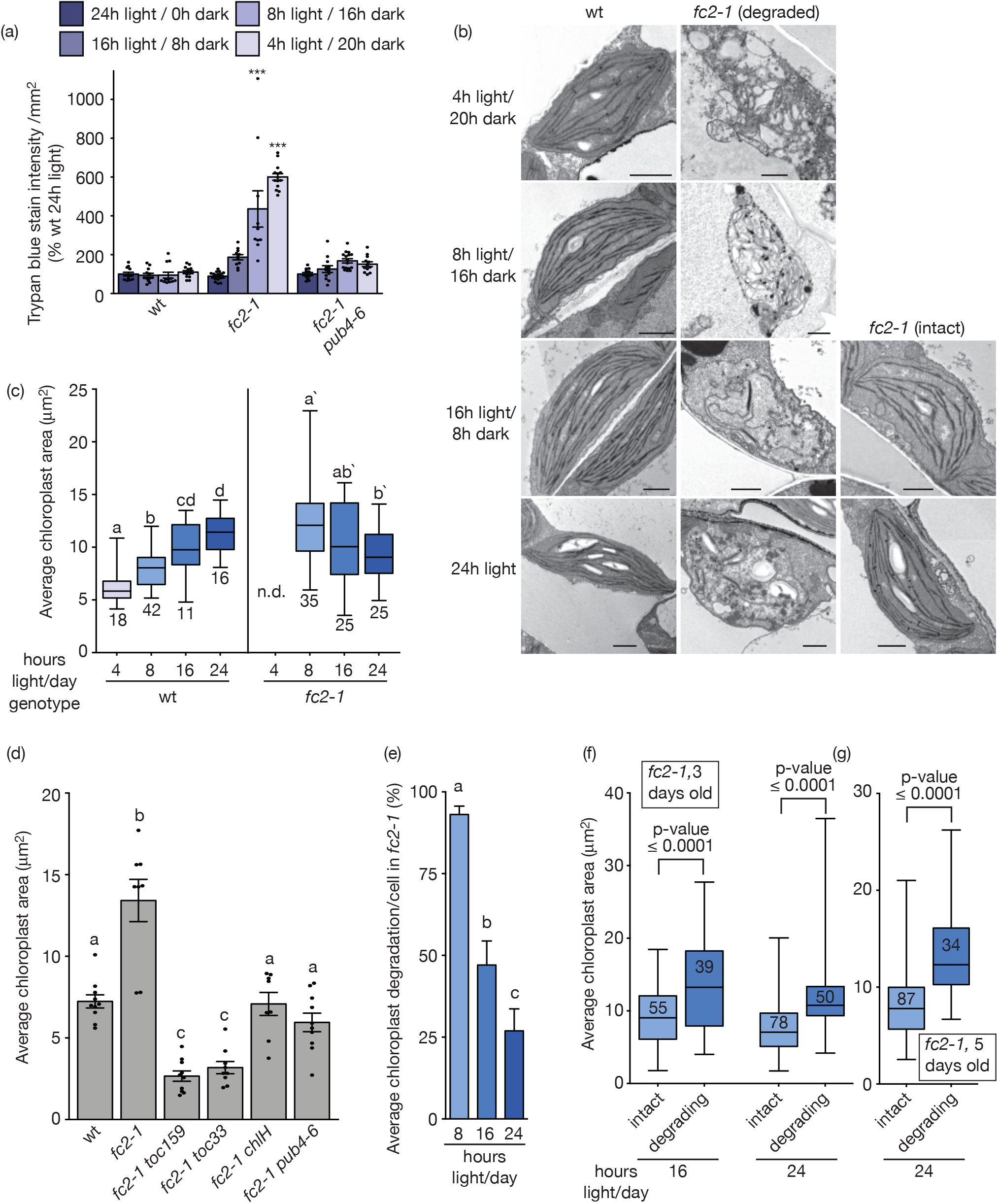
Chloroplast swelling is associated with photo-oxidative stress and degradation. The effect of the photoperiod on chloroplast degradation was assessed. **A)** Quantification of the trypan blue staining intensity/mm^2^ (% wt in 24h constant light) of six-day-old seedlings grown under the indicated photoperiods. Shown are means of biological replicates (n ≥ 10 seedlings) ± SEM. Representative images are shown in Fig. S4. **B)** Representative transmission electron microscopy (TEM) images of three-day-old (two hours post subjective dawn) wt and *fc2-1* chloroplasts in same photoperiods. Scale bars = 1 µm. **C)** Shown are box and whisker plots of mean chloroplast areas per cell (µm^2^, min to max) in seedlings grown under the indicated photoperiods. Chloroplast area in *fc2-1* seedlings grown under 4h light/20h dark cycling photoperiods was not determined (nd) due to extensive cellular degradation. Total number of cells assessed is indicated under whiskers. **D)** Shown are the mean chloroplast areas (µm^2^) in three-day-old seedlings grown under 6h light/18h dark diurnal cycling conditions (six hours post subjective dawn) ± SEM (n ≥ 8 chloroplasts from separate cells). **E)** Shown is the mean % chloroplast degradation per cell in three-day-old *fc2-1* seedlings (two hours post subjective dawn) grown under the indicated photoperiods (n ≥ 22 cells) ± SEM. **F and G)** *fc2-1* chloroplasts from panel C and Fig. 4b were divided into intact and degraded based on their morphological characteristics and measured separately. Shown are box and whisker plots of intact and degrading chloroplast areas (min to max) in three-day-old and five-day-old *fc2-1* seedlings grown under the indicated photoperiods (two hours post subjective dawn). Total number of chloroplasts measured indicated in boxes. All data shown in panels C-G was generated using TEM images. Statistical analyses in panels A, C-E were performed by one-way ANOVA tests followed by a Dunnett’s multiple comparisons test with the wild type (wt) 24h samples (A) or a Tukey-Kramer post-test (C-E). ***, p-value ≥ 0.001. Different letters above bar graph represent significant differences (p-value ≤ 0.05). In panel C, separate statistical analyses were performed on wt and *fc2-1* and the significance for *fc2-1* chloroplast areas is denoted by letters with a prime symbol (ʹ). Statistical analyses in F and G were performed by student’s t-tests. Closed circles represent individual data points.

Next, we assessed chloroplast ultrastructure in three-day-old seedlings grown under the same photoperiods (Fig. 5b). Seedlings were fixed for TEM two hours after subjective dawn to allow for the accumulation of ^1^O_2_ and the initiation of cellular degradation in the *fc2-1* mutant (Woodson et al. 2015). For wt chloroplasts, there was a clear positive correlation between average chloroplast area and length of day; from 6.08 µm^2^ under 4h light/20h dark conditions to 11.28 µm^2^ under 24h constant light conditions) (Fig. 5c). This light-dependent increase in size is likely due to longer days providing more light energy for chloroplast biogenesis. In *fc2-1* mutants, we observed the opposite pattern. Chloroplasts became larger under shorter days (9.34 µm^2^ under 24h constant conditions to 12.27 µm^2^ under 8h light/16h dark conditions) (Fig. 5c). We were unable to assess chloroplast size under 4h light/20h dark conditions as there was too much cellular degradation to accurately measure areas (Fig. 5b).

To determine if the swelling in *fc2-1* chloroplasts was ^1^O_2_-signaling dependent, we returned to a previous TEM data set of three-day-old seedlings grown under 6h light/18h dark diurnal cycling conditions and fixed six hours after subjective dawn (Woodson et al. 2015) (Fig. S2c). This set contained four genetic suppressors of *fc2-1* cell death and chloroplast degradation; *toc159* and *toc33* affecting the plastid protein import machinery, *chlH/gun5* affecting the tetrapyrrole pathway, and *pub4-6*. *toc159*, *toc33*, and *chlH* likely rescue *fc2-1* from cellular degradation by reducing ^1^O_2_ accumulation while *pub4-6* blocks ^1^O_2_ signaling (Woodson et al. 2015). As shown in Fig. 5d, each suppressor mutation reduces the average chloroplast area in the *fc2-1* background under these conditions. This suggests the observed chloroplast swelling in the *fc2-1* mutant is due to ^1^O_2_ accumulation and signaling.

Next, we wanted to test if chloroplast swelling was correlated to degradation within the same seedling. With the TEM images of three-day-old *fc2-1* seedlings used to measure chloroplast areas, we identified degrading chloroplasts by their irregular shapes, highly condensed/compressed grana, and few to non-existent thylakoids (Fig 5b). As shown in Fig. 5e, the ratio of intact and degrading chloroplasts changes with the photoperiod. When grown under 8h light/16h dark cycling light conditions, *fc2-1* chloroplasts were almost uniformly degraded (93%), consistent with cell death under these conditions. However, under 16h light/8h dark and 24 constant light conditions where significant cell death does not occur, 27% and 47% of chloroplasts remain intact, respectively. We hypothesized that there may be a difference in size and swelling between these two populations of chloroplasts and separately measured their areas (Fig. 5f). Under both 16h light/8h dark diurnal light cycling and 24h constant light conditions, degrading chloroplasts were significantly larger compared to intact chloroplasts (increases of 49% and 61% in cycling and constant light, respectively). Measuring chloroplast areas from TEM images of five-day-old *fc2-1* seedlings grown in 24h constant light showed that this difference was sustained in older seedlings (58% increase in size) (Fig 5g). Together, these results show a clear correlation between chloroplast swelling and degradation in the *fc2-1* mutant.

### Chloroplast degradation is associated with an increase in plastoglobule size

Plastids of all types contain globular lipid droplets called PGs that harbor proteins, most of which are involved in plastid transitions and lipid metabolism (Rottet et al. 2015; van Wijk and Kessler 2017). In addition to chloroplast swelling, PG size has also been shown to increase during chloroplast stress and senescence. To determine if changes in PG size is also associated with ^1^O_2_-induced chloroplast degradation, we measured PG size in the five-day-old wt, *pub4-6*, *fc2-1*, *fc2-2*, and *fc2-1 pub4-6* lines grown under 24h constant light using our TEM images (Fig. 6a). Under this condition, both *fc2-1* and *fc2-1 pub4-6* seedlings had significant increases in PG size (20% and 26%, respectively) compared to wt. Next, we measured the effect of the 24h photoperiod on PG size in three-day-old seedlings via TEM analysis. In wt seedlings, PG size was essentially unchanged in the four tested photoperiods (Fig. 6b). In *fc2-1* seedlings, PG size was also very similar in seedlings grown in 24h constant light or 16h light/8h dark cycles. In the 8h light/ 16h dark photoperiod however, there was a significant ∼3-fold increase in PG size. PGs could not be accurately discerned in chloroplasts grown in 4h light/20h dark, as the chloroplasts themselves had been completely degraded.

**Figure 6.**
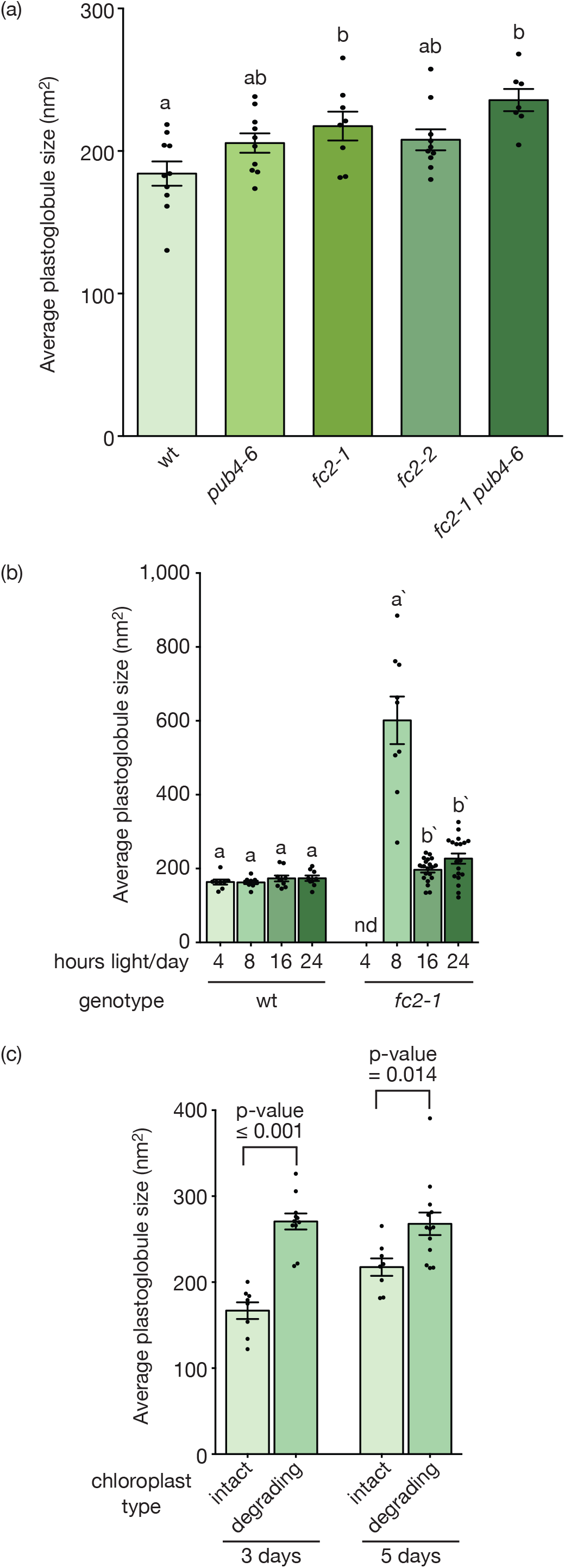
Chloroplast degradation is associated with an increase in plastoglobule size. Plastoglobule (PG) size was assessed in mutant seedlings grown in different photoperiods. Shown are the mean PG sizes (nm^2^ ± SEM) per chloroplast **A)** in five-day-old seedlings (n ≥ 7 chloroplasts) grown in 24h constant light and **B)** in three-day-old wt and *fc2-1* seedlings (n ≥ 8 chloroplasts) grown under the indicated photoperiods (two hours post subjective dawn). Data calculated from TEM images. **C**) PGs from intact and degrading chloroplasts were measured separately in three-and five-day-old *fc2-1* seedlings grown in constant light (n ≥ 8 chloroplasts). Shown are the mean PG sizes (nm^2^ ± SEM) per chloroplast. Statistical analyses were performed in A and B by one-way ANOVA and different letters above bars indicate significant differences determined by Tukey-Kramer post-tests (p value ≤ 0.05). In panel B, separate statistical analyses were performed for the wt and *fc2-1* seedlings and the significance for the *fc2-1* seedlings is also denoted with a prime symbol (ʹ). In panel C, the statistical tests were performed with a student’s t-test. In all bar graphs, closed circles represent individual data points. nd, not determined.

Together, these results suggested that an increase of PG size was associated with degrading chloroplasts in the *fc2-1* mutant. To further test this hypothesis, we separately analyzed intact and degrading chloroplasts in *fc2-1* seedlings grown under constant 24h light conditions. In both three-day-old and five-day-old seedlings, there was a significant increase in PG size in degrading chloroplasts (53% and 20% increase, respectively) (Fig. 6c). Together, these results suggest large PGs are indicators of ^1^O_2_ stress and degradation, and along with chloroplast swelling, precede cell death in the *fc2-1* mutant.

### Plastoglobule association with grana stacks precedes chloroplast degradation

PGs are mostly localized to the highly curved thylakoid margins within chloroplasts (Austin et al. 2006). However, under photo-oxidative stress, PGs also associate with the thylakoid grana stacks, which has been shown to be an indicator of ^1^O_2_ stress (Wang et al. 2020). To determine if PGs were behaving this way under our growth conditions, we first calculated the percentage of PGs on the grana (defined as PGs spanning two or more thylakoids (Fig. S5)) in TEM images of five-day-old seedlings grown in constant light (Fig. 7a). As this calculation is dependent on chloroplasts having normal internal membrane structures, only intact chloroplasts were considered. Under these conditions, both *fc2-1* and *fc2-2* chloroplasts had about twice as many PGs on the grana compared to wt. This increase of grana-associated PGs in the *fc2-1* mutant was suppressed by the *pub4-6* mutation suggesting such structures are dependent on ^1^O_2_ signaling.

**Figure 7.**
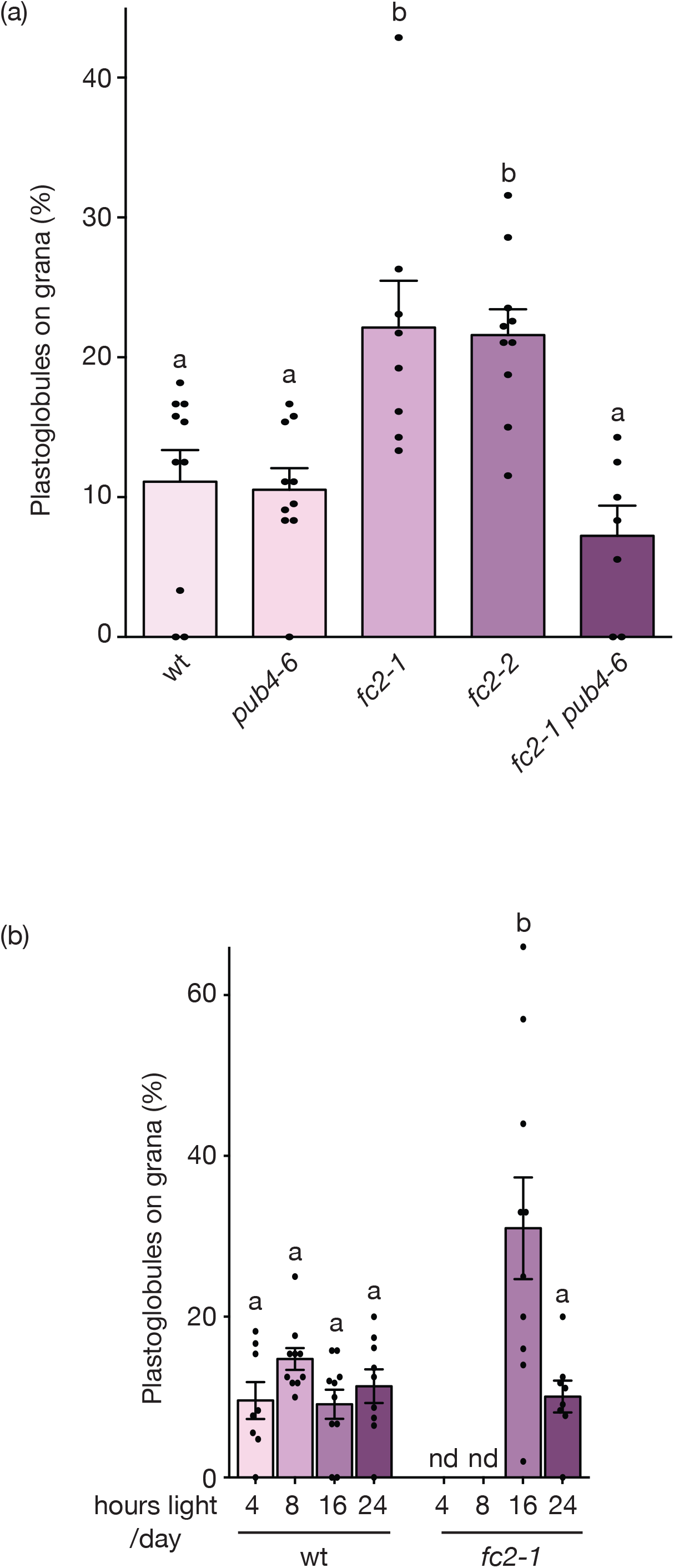
Plastoglobuli associate with grana stacks in *fc2* mutants. The portion of plastoglobuli (PGs) associated with grana was assessed in mutant seedlings grown in different photoperiods. **(A)** Shown are the average percentage of PGs (±SEM) on the grana in five-day-old seedling chloroplasts (n ≥ 7 chloroplasts). **(B)** Shown are the average percentage of PGs (± SEM) on the grana in three-day-old seedlings grown under the indicated photoperiods (n ≥ 8 chloroplasts). Calculations based on TEM images of chloroplasts. Statistical analyses were performed by one-way ANOVA and different letters above bars indicate significant differences determined by Tukey-Kramer post-tests (p value ≤ 0.05). In all bar graphs, closed circles represent individual data points. nd, not determined.

Next, we analyzed the association of PGs on the grana in three-day-old wt and *fc2-1* chloroplasts under different photoperiods using our TEM images. As grana are mostly absent in degrading chloroplasts, only intact chloroplasts from 16h light/8h dark cycling and 24h constant light conditions could be used in the analysis. In wt, the photoperiod had no significant effect on PG/grana association (Fig. 7b). In *fc2-1* chloroplasts, however, PG association with the grana was increased under 16h light/8h dark cycling light compared to 24h constant light. Together, these results suggest the association of PGs with grana may be an early indicator of ^1^O_2_ stress occurring prior to swelling, degradation, or interactions with the central vacuole.

### Chloroplast senescence is initiated prematurely in *fc2* mutant seedlings

In addition to photo-oxidative stress, PGs are also known to play a role in leaf senescence and chloroplast disassembly by harboring enzymes involved in chlorophyll breakdown and lipid remodeling (Rottet et al. 2015; van Wijk and Kessler 2017). To determine if such pathways may be activated prematurely in *fc2-1* mutants, we monitored the expression of two such PG marker genes involved in chloroplast disassembly. *PHEOPHYTIN PHEOPHORBIDE HYDROLASE* (*PPH*) encodes an enzyme that releases the phytol group from chlorophyll during catabolism (Schelbert et al. 2009). *PHYTYL EASTER SYNTHASE 1* (*PES1*) encodes a diacylglycerol acetyltransferase that can esterify phytol with fatty acids released from thylakoid galactolipids to form triacylglycerol and phytyl fatty acid esters. Such metabolites contribute to the large PGs found in senescing chloroplasts (gerontoplasts) (Lippold et al. 2012). Both enzymes are also associated with PGs (Lundquist et al. 2012; Ytterberg et al. 2006) and may serve as markers for chloroplast senescence.

An analysis of *PPH* and *PES1* steady state transcripts showed that both are significantly up-regulated in the *fc2-1* mutant compared to wt (Fig. 8). This was observed in both 24h constant and 6h light/18h dark cycling light conditions suggesting that these genes were not merely responding to general cellular degradation or cell death in the *fc2-1* mutant. Therefore, in addition to abiotic stress signaling, *fc2-1* chloroplasts may also be initiating a senescence pathway for their own dismantling and recycling.

**Figure 8.**
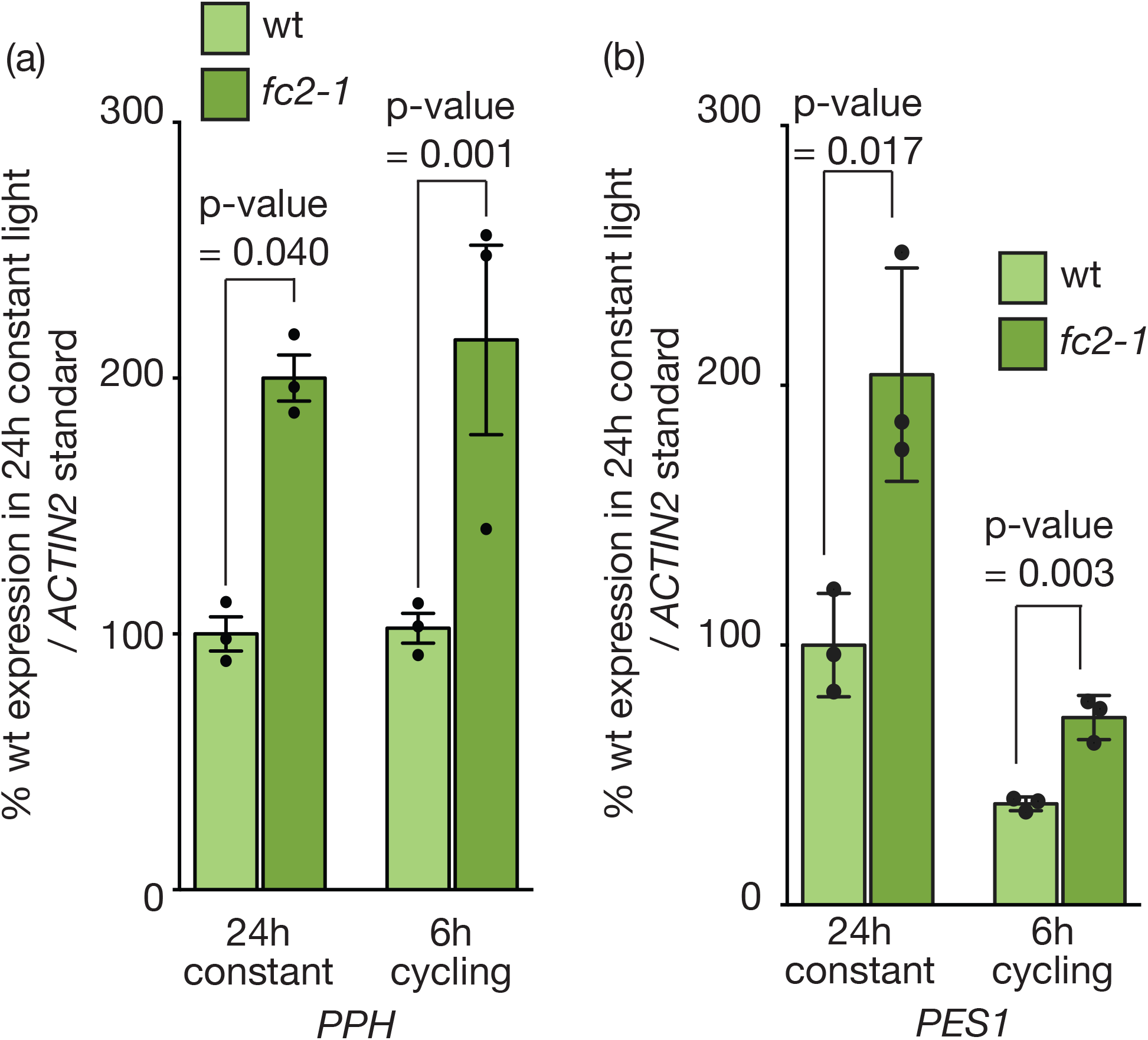
Plastoglobule stress genes are induced in the *fc2-1* mutant. The expression of two plastoglobule (PG) stress marker genes, *PPH* and *PES1*, were measured in the *fc2-1* mutant. Shown are mean levels of **A)** *PPH* and **B)** *PES1* steady-state transcript levels from four-day-old seedlings grown in constant (24h) or 6h light/18h dark (6h) cycling light as measured by RT-qPCR (n = 3 biological replicates) ± SEM. Values were normalized to *ACTIN2* expression. Statistical analysis was performed by student’s t-tests. p-values are denoted above. Closed circles represent individual data points.

## Discussion

In plant cells, chloroplast degradation has been demonstrated to be induced by photo-oxidative damage and starvation (Zhuang and Jiang 2019). However, little is known about the process of degradation, how chloroplasts interact with the central vacuole, or how multiple degradation pathways may interact. As such, a better understanding of the associated structures may offer much needed insight into the mechanisms of chloroplast degradation and how individual chloroplasts are selected for turnover. To monitor this phenomenon, here we have used the Arabidopsis *fc2* mutant that has accelerated chloroplast turnover due to the accumulation of ^1^O_2_ in the chloroplasts. Under short day diurnal light cycling conditions, *fc2* mutants suffer from wholesale chloroplast degradation and eventual cell death due to high levels of Proto (1620 pmoles/g fresh weight (Woodson et al. 2015)) and ^1^O_2_ (2.5-fold increase compared to wt (Alamdari et al. 2021)). However, under permissive 24h constant light conditions, only a subset of chloroplasts is selected for degradation and plants avoid cell death. Under these conditions, we demonstrated Proto and ^1^O_2_ levels are still increased in *fc2* mutants, but to a lesser degree (Table 1 and Fig. S1). This indicates *fc2* mutants still experience mild photo-oxidative stress under 24h constant light conditions, but below a threshold level for cell death. Therefore, such permissive conditions allow for the study of selective chloroplast degradation in otherwise healthy cells.

When *fc2* mutants are grown under 24h constant light conditions, degrading chloroplasts can be easily identified by their irregular shapes, compressed grana, disorganized internal membranes, and unusual vesicles (Alamdari et al. 2020; Alamdari et al. 2021; Lemke et al. 2021; Woodson et al. 2015). As previously reported, a subset of these damaged chloroplasts appears to protrude into the central vacuole with “blebbing” structures (Fig. 1a) (Woodson et al. 2015). A closer look at these structures suggests they are formed by close association of the chloroplast envelope and tonoplast membranes (Fig. 1b). Other proximal cytoplasmic structures, such as mitochondria, appear unaffected (Fig. 1c) suggesting the chloroplast is being specifically targeted for turnover in the central vacuole. The lack of any additional membranes further supports the hypothesis that autophagosomes are not involved in these structures and the process resembles a fission-type microautophagy pathway independent of the core autophagy machinery (Lemke et al. 2021).

A SBF-SEM 3D analysis of these structures offered additional important conclusions. First, these measurements confirmed that degrading chloroplasts remained in the cytoplasm whilst the blebbing structure itself was entirely located within the central vacuole. Second, these structures were much more common than the 2D TEM imaging indicated. In our 2D tile-scanned FE-SEM data, such structures were identified in > 0.5% of *fc2* chloroplasts (Fig. 1f). In the 3D volumetric SBF-SEM datasets, almost 8% of chloroplasts were associating with blebbing vacuole structures. This difference is likely due to the small contact point between the degrading chloroplast and blebbing structure (∼7 µm^2^), which would make capturing such associations difficult when assessing traditional 2D TEM (our samples were only 70-100 nm thick). Third, the blebbing structures were variable in size. Most were small (9 µm^2^, 14% of the chloroplast volume), but we observed one structure of considerable size (178 µm, 202% of the chloroplast volume) (Fig. S3a-d). It is not clear if this is a different class of structure or a different time point in the process. Interestingly, the associated chloroplast appears normal suggesting it may represent an early stage of chloroplast degradation. Finally, such structures were completely absent in *fc2-1 pub4-6* mutants, confirming ^1^O_2_ signals are necessary for their induction.

The above work suggests chloroplast/vacuole interactions are much more frequent than first assumed, but it is still unclear why certain chloroplasts are targeted for degradation. To address this, we used a previously published FE-SEM data set to measure chloroplast degradation in entire cotyledon cross sections of *fc2-1* cells (Woodson et al. 2015). The position of the chloroplast within the cell (adaxial vs. abaxial) did not significantly influence the chance of degradation (Fig. 3b). This was somewhat surprising as we hypothesized that a closer proximity to light on the adaxial side of the cell could serve to increase ROS production. The thin size of seedling cotyledons and light scattering in mesophyll cells, however, may make such positions negligible.

Chloroplast degradation was more common in the abaxial-localized spongy mesophyll cells (at least when only cells with degrading chloroplast were considered) (Fig. 3a). The reason for this difference is not clear. The two cell types are morphologically distinct and play different roles in leaf physiology, but their molecular differences are not well-defined (Pyke and López-Juez 1999). Therefore, it is possible chloroplasts in spongy mesophyll cells are more susceptible to photo-oxidative damage and/or more likely to be degraded. Being in the lower layer of cells, the photochemistry of spongy mesophyll cells may be tuned to lower light intensities than palisade cells (Terashima and Inoue 1984). Alternatively, the *fc2-1* mutation may have different effects in the two cell types, leading to different levels of tetrapyrroles and ^1^O_2_ production. One study has shown at least one transcript encoding a tetrapyrrole biosynthetic enzyme (protochlorophyllide oxidoreductase) is differentially expressed between these two cell types (Marrison et al. 1996).

The observation that single chloroplasts can be removed from cells is indicative of a mechanism by which a cell can differentiate between healthy and severely damaged chloroplasts. One possible feature that may allow for recognition is organelle swelling, which has been shown to occur prior to chlorophagy during excess light stress (Nakamura et al. 2018). In that case, overexpression of *VESICLE-INDUCING PROTEIN IN PLASTIDS 1* (*VIPP1*) suppressed chloroplast swelling and movement to the vacuole. As this protein has been demonstrated to regulate chloroplast envelope integrity (Zhang et al. 2012), this result suggested swelling may be a mechanism by which damaged chloroplasts are identified. Here, we also show there is a strong correlation between chloroplast swelling and degradation in the *fc2* mutants. First, we see an inverse relationship between day length and chloroplast size in the *fc2-1* mutant. An increase in photo-oxidative stress and cell death in shorter days is associated with an increase in chloroplast degradation and swelling (Figs. 5a and c). Second, we observed a difference in size between intact and degrading chloroplasts within the same cells of healthy seedlings grown under permissive 24h light conditions (Figs. 5f and g). Finally, chloroplast swelling is dependent on chloroplast ^1^O_2_ signals, as it is blocked by the four tested *fts* mutations (Fig. 5d).

Together, these results are consistent with chloroplast swelling preceding (and possibly initiating) degradation in *fc2* mutants. Therefore, chloroplast swelling may well be a general mechanism to indicate organelle dysfunction and to promote turnover. While chloroplast swelling is also involved in chlorophagy, it may be expected that the cellular machinery involved in recognizing damaged chloroplasts and initiating interactions with the central vacuole are specific and distinct. Previous genetic analyses of ^1^O_2_-induced chloroplast degradation in the *fc2* mutant and chlorophagy have suggested different pathways are involved. Chlorophagy is not dependent on PUB4 (Kikuchi et al. 2020), and chloroplast blebbing in *fc2* mutants can occur in the absence of the core autophagy machinery (Lemke et al. 2021). There are also several structural differences between the chloroplasts targeted by chlorophagy (relatively intact before being transported to the central vacuole) and ^1^O_2_-dependent degradation (degrade within the cytoplasm prior to interacting the central vacuole). Finally, chlorophagy after UV-B stress appears to be induced by superoxide/hydrogen peroxide, rather than by ^1^O_2_.

Our results also demonstrated PGs are highly dynamic in *fc2* mutants, particularly when stressed. First, PG size increased under 8h light/16h dark diurnal cycling conditions (Fig. 6b), indicating a high degree of photo-oxidative stress. This may indicate PGs are aiding in thylakoid protection or disassembly, similar to what has been observed during photo-oxidative stress or senescence (Austin et al. 2006; Besagni and Kessler 2013). Second, within the same cells under 24h constant light conditions, degrading chloroplasts contained larger PGs than healthy chloroplasts (Fig. 6c). Such an effect also appears to be cumulative as the difference was more pronounced in five-day-old seedlings compared to three-day-old seedlings (Fig. 6c).

PGs were also more associated with grana in the *fc2-1* mutant, particularly under light cycling conditions (Fig. 7b). Such an association has previously been demonstrated to occur in the *safeguard1* mutant that suffers from chloroplast ^1^O_2_ stress (Wang et al. 2020). This may indicate grana are among the first targets of oxidative damage and a protective mechanism is being induced. PGs also contain *α*-tocopherol, which plays a role in protecting thylakoids and their proteins from oxidation (Eugeni Piller et al. 2014). Interestingly, the *pub4-6* mutation reversed the association of PGs associating with grana, but did not reverse the enlargement of PGs (Figs. 6a and 7a). This suggests the enlargement of PGs is not necessarily a consequence of degrading chloroplasts, but may be an indicator of stress and an increase of lipoprotein and antioxidant production (Rottet et al. 2015; van Wijk and Kessler 2017). Conversely, increased association of PGs with the grana precedes chloroplast degradation and is dependent on ^1^O_2_ signals. As such, it may be an indicator of chloroplast degradation rather than a protective mechanism.

In addition to stress, increased PG size is also associated with senescence (Ghosh et al. 2001). To test if such a pathway is activated in *fc2* mutants and involved in chloroplast degradation, we monitored the expression of two genes encoding PG enzymes involved in senescence and chloroplast disassembly; *PPH* and *PES1* (Figs. 8a and b). Compared to wt, both genes were upregulated in *fc2-1* mutants in healthy and degrading cells. This indicated PGs may have a disassembly role in *fc2* mutants and a senescence pathway has been activated. Previous work on chloroplast degradation has found other parallels with senescence. Canonical autophagosomes are not only involved in chlorophagy during stress (Izumi et al. 2017; Nakamura et al. 2018), but also non-specific turnover of chloroplasts during dark-induced carbon starvation and senescence (Wada et al. 2009). In rice, the CV pathway has been shown to be important in regulating photo-assimilation of nitrogen by controlling chloroplast turnover during drought-induced senescence (Sade et al. 2018). *pub4-6* mutants also have accelerated senescence phenotypes, particularly in backgrounds defective in the core autophagy machinery (Kikuchi et al. 2020; Woodson et al. 2015). As ROS generally accumulates in senescing leaves (Juvany et al. 2013), it may also be able to induce non-selective chloroplast degradation under such conditions. Thus, ^1^O_2_-induced chloroplast degradation may play a dual role in redistributing nutrients while also protecting cells from photo-oxidative damage.

## Conclusions

Our analysis of degrading chloroplasts in the *fc2* mutant has revealed several important observations about the process of ^1^O_2_-damaged chloroplast turnover. First, a third of such degrading chloroplasts in *fc2* cells are in some state of interaction with the central vacuole, where they protrude with structures resembling a fission-type microautophagy process. In addition to the disorganized internal membranes, degrading chloroplasts were also shown to swell and contain large PGs. Except for PG enlargement, these features are dependent on chloroplast ^1^O_2_ signals and point to the possibility that chloroplast swelling is possible mechanism for recognition in the cell. Moreover, the dynamic PGs in *fc2* chloroplasts suggest a connection between ^1^O_2_-chloroplast degradation and senescence, indicating the cell is balancing stress tolerance and nutrient availability. Future studies should focus on how such chloroplast structures are recognized by the cell for turnover, the overlap between stress and senescence, and the possible interactions between degradation pathways. A deeper understanding of the mechanisms involved in chloroplast turnover should provide additional tools to develop plants and crops that cope with dynamic environments by balancing stress with cellular degradation.

## Materials and Methods

### Biological material, growth conditions, and treatments

In this study, the wt control line used in all experiments was *Arabidopsis thaliana* ecotype *Columbia* (Col-0). Table S1 lists the mutant lines used. Sequence data for the T-DNA lines came from the GABI (Kleinboelting et al. 2012) and SAIL (Sessions et al. 2002) collections; *fc2-1* (GABI_766H08) and *fc2-2* (SAIL_20_C06C) were described previously (Woodson et al. 2011). Lines *pub4-6*, *fc2-1 pub4-6*, *fc2-1 toc159*, *fc2-1 toc33*, and *fc2-1 chlH* were also described previously (Woodson et al. 2015).

Seeds were surface sterilized using 30% liquid bleach (v:v) with 0.04% Triton X-100 (v:v) for ten minutes as previously described (Alamdari et al. 2020). A three-four day cold treatment of 4°C in the dark was used to stratify seeds before transferring them to controlled artificial growth conditions of constant light or diurnal light/dark cycling conditions of approximately ∼100 μmol m^-2^ sec^-1^ white light using cool white fluorescent bulbs. Plants were grown at a constant temperature of 22°C.

### Analysis of FE-SEM data; quantification of chloroplast degradation and blebbing

To quantify chloroplast degradation, blebbing, and association with the central vacuole, a previously generated data set of FE-SEM images of cotyledon cross-sections were used (Woodson et al. 2015). For each genotype, two independent cross sections made from separate seedlings were available. To determine if chloroplast positioning within a cell or cell type within the cotyledon tissue influences the likelihood of degradation, 75 mesophyll cells (39 palisade cells and 36 mesophyll cells) from the two *fc2-1* seedlings were analyzed. Cells were first assigned to either the palisade or spongy layer category based upon position, size, and arrangement, and the total number of chloroplasts in each cell was counted. Degrading chloroplasts were noted based on their unique characteristics (compressed and disorganized internal membranes) and the percent degrading chloroplasts were calculated per cell. The percentage of degrading chloroplasts in each cell type was calculated for the overall population and for the sub-population of cells that contained at least one degrading chloroplast (20 palisade and 15 spongy mesophyll cells).

To determine if chloroplast position within a cell influences the chances of degradation, each cell was split along the equator into a “top” and “bottom” half and the percentage of degrading chloroplasts was calculated from the total number of chloroplasts in each half. Similarly, the overall percentage of degrading chloroplasts in the top versus bottom half of the cell for the total population and for the cells with at least one degrading chloroplast were calculated. Blebbing chloroplasts were identified in mesophyll cells by their association with large vacuolar protrusions.

### Transmission Electron microscopy

Three- and five-day-old seedlings were fixed and imaged by TEM as previously described (Woodson et al. 2015).

### Starch granule, chloroplast size, plastoglobule size, and plastoglobule positioning measurements

Transmission electron micrographs of three- or five-day-old seedlings were analyzed to calculate starch granule size, chloroplast area, % chloroplast degradation, PG size, and PG association with thylakoid grana stacks. For starch granules, the areas of individual chloroplasts and starch granules were measured using ImageJ software. If multiple starch granules were present, the individual areas were combined. An average chloroplast area and starch granule area was then calculated for each cell (at least three chloroplasts per cell), and starch granule area was then determined as a percentage of average chloroplast area from the combined averages (n ≥ 7 cells). Using ImageJ software, average chloroplast area was calculated by measuring the planar area of individual chloroplasts in at least 7 cells from each genotype and condition. For three-day-old wt seedlings grown in 16h light/8h dark diurnal light cycling conditions and all three-day-old seedlings grown under 6h light/18h dark diurnal light cycling conditions, one representative chloroplast per cell was measured. For all other samples, 2-12 chloroplasts were averaged per cell. For *fc2-1* cells grown in 16h light/8h dark cycling and 24h constant light conditions, chloroplasts were also separated into “intact” and “degrading” based on morphological characteristics. Degrading chloroplasts were identified by their disorganized internal membrane structures and irregular shapes and % chloroplast degradation was assessed on a per cell basis (n ≥ 22 cells). For PG size, chloroplasts (n ≥ 7) were selected from at least 7 separate cells and the average area of PGs within each chloroplast were measured using ImageJ. For PG association with grana stacks, the percentage of PGs on the grana was calculated from the same chloroplast images used to calculate PG size. PGs were considered to be on the grana if they were observed spanning at least two thylakoids (Fig. S5). In all cases, only chloroplasts from cotyledon mesophyll cells were considered.

### SBF-SEM analysis

For three dimensional SBF-SEM analysis, five-day-old wt and *fc2-1* seedlings grown in 24h constant light were used. An in-situ microtome (3-View, Gatan, Pleasanton, CA) housed within a variable pressure Scanning Electron Microscope, or SEM (Sigma VP, Zeiss), was used to sequentially take a backscatter image, slice 70 nm into the tissue, and capture another backscatter image again. This was performed at an image size of 5,120 pixels by 5,120 pixels over 750 slices, resulting in a total volume of 88 µm x 88 µm x 52.5 µm at a resolution of 17 nm/px in x and y, and 70 nm/px (the slice thickness) in the z-direction. To minimize sample charging due to the large voids of unstained tissue in the central vacuole, the SEM was operated at 1.75 KeV in variable pressure mode with a chamber pressure of 7 Pa.

Individual images acquired from the SBF-SEM analysis were opened in Fiji using the Bio-Formats plugin (Linkert et al. 2010). They were then concatenated to create an image stack which was further corrected for drift with the "StackReg" plugin in Fiji (Thévenaz et al. 1998). To count and assess chloroplasts, images were aligned in Image J and cells were identified (5 and 13 cells in wt and *fc2-1*, respectively). Following this, individual “chloroplast-blebbing” events were identified in *fc2-1* datasets. ROIs of these events were created to create smaller image stacks. These ROIs were then imported into Imaris 9.6 and chloroplasts and blebs were manually segmented using the "Contour" tool to create 3D surface volumes. Snapshots and 3D-renderings of these surfaces were then created with Imaris 9.6 using the "Snapshot" and "Animation" tools, respectively.

### RNA extraction and Real-time quantitative PCR

The RNeasy Plant Mini Kit (Qiagen) was used to extract total RNA from whole seedlings. Next, cDNA was synthesized using the Maxima first strand cDNA synthesis kit for RT-qPCR with DNase (Thermo Scientific) following the manufacturer’s instructions. Real-time PCR was performed using the SYBR Green Master Mix (BioRad) with the SYBR Green fluorophore and a CFX Connect Real Time PCR Detection System (BioRad). The following 2-step thermal profile was used in all RT-qPCR: 95 °C for 3 min, 40 cycles of 95 °C for 10s and 60 °C for 30s. *Actin2* expression was used as a standard to normalize all gene expression data. Table S2 lists the primers used.

### Trypan Blue staining

Trypan blue staining was used to assess cell death in leaves and cotyledons using a previously described method (Woodson et al. 2015). To quantify cell death, trypan blue intensity was measured with ImageJ using at least ten seedlings from each genotype and condition. Seedlings from each photoperiod were imaged separately under similar lighting conditions. To account for any small lighting differences, trypan blue signal was calculated by subtracting background intensity in each image.

### Monitoring singlet oxygen accumulation

Singlet Oxygen Sensor Green (SOSG, Molecular Probes) was used to assess bulk in vivo levels of ^1^O_2_ in four-day-old seedlings as previously described (Alamdari et al. 2020).

### HPLC measurements of tetrapyrrole intermediates

The levels of the five tetrapyrrole intermediates; Protoporphyrin IX (Proto), Uroporphyrinogen III (Uro III), Coproporphyrinogen III (Copro III), Mg-Protoporphyrin IX (Mg-Proto), and Mg-Protoporphyrin IX monomethylester (Mg-Proto-Me) were extracted from shoot tissue of three-day-old seedlings and measured as previously described (Woodson et al. 2015), using an adaptation of an HPLC/fluorescence detection protocol (Czarnecki et al. 2011).

## Supporting information

Fisher et al. 2021 supporting information

## Acknowledgements

The authors thank Matthew Lemke (University of Arizona) for technical assistance with trypan blue stains and Kamran Alamdari (University of Arizona) for technical assistance with RT-qPCR.

## References

Alamdari, K., Fisher, K.E., Sinson, A.B., Chory, J. and Woodson, J.D. (2020) Roles for the chloroplast-localized PPR Protein 30 and the "Mitochondrial" Transcription Termination Factor 9 in chloroplast quality control. Plant J 103: 735–751.

Alamdari, K., Fisher, K.E., Tano, D.W., Rai, S., Palos, K.R., Nelson, A.D.L., et al. (2021) Chloroplast quality control pathways are dependent on plastid DNA synthesis and nucleotides provided by cytidine triphosphate synthase two. New Phytol.

Asada, K. (2006) Production and scavenging of reactive oxygen species in chloroplasts and their functions. Plant Physiol 141: 391–396.

Austin, J.R., Frost, E., Vidi, P.A., Kessler, F. and Staehelin, L.A. (2006) Plastoglobules are lipoprotein subcompartments of the chloroplast that are permanently coupled to thylakoid membranes and contain biosynthetic enzymes. Plant Cell 18: 1693–1703.

Besagni, C. and Kessler, F. (2013) A mechanism implicating plastoglobules in thylakoid disassembly during senescence and nitrogen starvation. Planta 237: 463–470.

Chan, K.X., Mabbitt, P.D., Phua, S.Y., Mueller, J.W., Nisar, N., Gigolashvili, T., et al. (2016) Sensing and signaling of oxidative stress in chloroplasts by inactivation of the SAL1 phosphoadenosine phosphatase. Proc Natl Acad Sci U S A 113: E4567–4576.

Chan, K.X., Phua, S.Y., Crisp, P., McQuinn, R. and Pogson, B.J. (2015) Learning the Languages of the Chloroplast: Retrograde Signaling and Beyond. Annu Rev Plant Biol 67: 25–53.

Chiba, A., Ishida, H., Nishizawa, N.K., Makino, A. and Mae, T. (2003) Exclusion of ribulose-1,5-bisphosphate carboxylase/oxygenase from chloroplasts by specific bodies in naturally senescing leaves of wheat. Plant Cell Physiol 44: 914–921.

Czarnecki, O., Peter, E. and Grimm, B. (2011) Methods for analysis of photosynthetic pigments and steady-state levels of intermediates of tetrapyrrole biosynthesis. Methods Mol Biol 775: 357–385.

de Souza, A., Wang, J.Z. and Dehesh, K. (2017) Retrograde Signals: Integrators of Interorganellar Communication and Orchestrators of Plant Development. Annu Rev Plant Biol 68: 85–108.

Dogra, V. and Kim, C. (2019) Singlet Oxygen Metabolism: From Genesis to Signaling. Front Plant Sci 10: 1640.

Eugeni Piller, L., Glauser, G., Kessler, F. and Besagni, C. (2014) Role of plastoglobules in metabolite repair in the tocopherol redox cycle. Front Plant Sci 5: 298–298.

Foyer, C.H. (2018) Reactive oxygen species, oxidative signaling and the regulation of photosynthesis. Environ Exp Bot 154: 134–142.

Ghosh, S., Mahoney, S.R., Penterman, J.N., Peirson, D. and Dumbroff, E.B. (2001) Ultrastructural and biochemical changes in chloroplasts during Brassica napus senescence. Plant Physiology and Biochemistry 39: 777–784.

Harnischfeger, G. (1973) Chloroplast Degradation in Ageing Cotyledons of Pumpkin. J Exp Bot 24: 1236–1244.

Izumi, M., Ishida, H., Nakamura, S. and Hidema, J. (2017) Entire Photodamaged Chloroplasts Are Transported to the Central Vacuole by Autophagy. Plant Cell 29: 377–394.

Juvany, M., Müller, M. and Munné-Bosch, S. (2013) Photo-oxidative stress in emerging and senescing leaves: a mirror image? J Exp Bot 64: 3087–3098.

Kikuchi, Y., Nakamura, S., Woodson, J.D., Ishida, H., Ling, Q., Hidema, J., et al. (2020) Chloroplast Autophagy and Ubiquitination Combine to Manage Oxidative Damage and Starvation Responses. Plant Physiol 183: 1531–1544.

Kim, I., Rodriguez-Enriquez, S. and Lemasters, J.J. (2007) Selective degradation of mitochondria by mitophagy. Archives of Biochemistry and Biophysics 462: 245–253.

Kleinboelting, N., Huep, G., Kloetgen, A., Viehoever, P. and Weisshaar, B. (2012) GABI-Kat SimpleSearch: new features of the Arabidopsis thaliana T-DNA mutant database. Nucleic Acids Res 40: D1211–1215.

Lemke, M.D., Fisher, E.M., Kozlowska, M.A., Tano, D.W. and Woodson, J.D. (2021) The core autophagy machinery is not required for chloroplast singlet oxygen-mediated cell death in the Arabidopsis plastid ferrochelatase two mutant. BMC Plant Biol??: ??

Linkert, M., Rueden, C.T., Allan, C., Burel, J.M., Moore, W., Patterson, A., et al. (2010) Metadata matters: access to image data in the real world. J Cell Biol 189: 777–782.

Lippold, F., vom Dorp, K., Abraham, M., Hölzl, G., Wewer, V., Yilmaz, J.L., et al. (2012) Fatty acid phytyl ester synthesis in chloroplasts of Arabidopsis. Plant Cell 24: 2001–2014.

Lu, Y. and Yao, J. (2018) Chloroplasts at the Crossroad of Photosynthesis, Pathogen Infection and Plant Defense. Int J Mol Sci 19: 3900.

Lundquist, P.K., Poliakov, A., Bhuiyan, N.H., Zybailov, B., Sun, Q. and van Wijk, K.J. (2012) The functional network of the Arabidopsis plastoglobule proteome based on quantitative proteomics and genome-wide coexpression analysis. Plant Physiol 158: 1172–1192.

Makino, A. and Osmond, B. (1991) Effects of Nitrogen Nutrition on Nitrogen Partitioning between Chloroplasts and Mitochondria in Pea and Wheat. Plant Physiol 96: 355–362.

Marrison, J.L., Schunmann, P., Ougham, H.J. and Leech, R.M. (1996) Subcellular Visualization of Gene Transcripts Encoding Key Proteins of the Chlorophyll Accumulation Process in Developing Chloroplasts. Plant Physiol 110: 1089–1096.

Martinez, D.E., Costa, M.L., Gomez, F.M., Otegui, M.S. and Guiamet, J.J. (2008) ’Senescence-associated vacuoles’ are involved in the degradation of chloroplast proteins in tobacco leaves. Plant J 56: 196–206.

Michaeli, S., Honig, A., Levanony, H., Peled-Zehavi, H. and Galili, G. (2014) Arabidopsis ATG8-INTERACTING PROTEIN1 Is Involved in Autophagy-Dependent Vesicular Trafficking of Plastid Proteins to the Vacuole. Plant Cell 26: 4084–4101.

Nakamura, S., Hidema, J., Sakamoto, W., Ishida, H. and Izumi, M. (2018) Selective Elimination of Membrane-Damaged Chloroplasts via Microautophagy. Plant Physiol 177: 1007–1026.

Nakamura, S. and Izumi, M. (2018) Regulation of Chlorophagy during Photoinhibition and Senescence: Lessons from Mitophagy. Plant Cell Physiol 59: 1135–1143.

Niwa, Y., Kato, T., Tabata, S., Seki, M., Kobayashi, M., Shinozaki, K., et al. (2004) Disposal of chloroplasts with abnormal function into the vacuole in Arabidopsis thaliana cotyledon cells. Protoplasma 223: 229–232.

Otegui, M.S., Noh, Y.S., Martinez, D.E., Vila Petroff, M.G., Staehelin, L.A., Amasino, R.M., et al. (2005) Senescence-associated vacuoles with intense proteolytic activity develop in leaves of Arabidopsis and soybean. Plant J 41: 831–844.

Pyke, K. and López-Juez, E. (1999) Cellular Differentiation and Leaf Morphogenesis in Arabdopsis. Critical Reviews in Plant Sciences 18: 527–546.

Pyke, K.A. and Leech, R.M. (1994) A Genetic Analysis of Chloroplast Division and Expansion in Arabidopsis thaliana. Plant Physiol 104: 201–207.

Rottet, S., Besagni, C. and Kessler, F. (2015) The role of plastoglobules in thylakoid lipid remodeling during plant development. Bba-Bioenergetics 1847: 889–899.

Sade, N., Umnajkitikorn, K., Rubio Wilhelmi, M.D.M., Wright, M., Wang, S. and Blumwald, E. (2018) Delaying chloroplast turnover increases water-deficit stress tolerance through the enhancement of nitrogen assimilation in rice. J Exp Bot 69: 867–878.

Scharfenberg, M., Mittermayr, L., E, V.O.N.R.-L., Schlicke, H., Grimm, B., Leister, D., et al. (2015) Functional characterization of the two ferrochelatases in Arabidopsis thaliana. Plant Cell Environ 38: 280–298.

Schelbert, S., Aubry, S., Burla, B., Agne, B., Kessler, F., Krupinska, K., et al. (2009) Pheophytin pheophorbide hydrolase (pheophytinase) is involved in chlorophyll breakdown during leaf senescence in Arabidopsis. Plant Cell 21: 767–785.

Sessions, A., Burke, E., Presting, G., Aux, G., McElver, J., Patton, D., et al. (2002) A high-throughput Arabidopsis reverse genetics system. Plant Cell 14: 2985–2994.

Suo, J., Zhao, Q., David, L., Chen, S. and Dai, S. (2017) Salinity Response in Chloroplasts: Insights from Gene Characterization. Int J Mol Sci 18: 1011.

Terashima, I. and Inoue, Y. (1984) Comparative Photosynthetic Properties of Palisade Tissue Chloroplasts and Spongy Tissue Chloroplasts of Camellia japonica L.: Functional Adjustment of the Photosynthetic Apparatus to Light Environment within a Leaf. Plant and Cell Physiology 25: 555–563.

Thévenaz, P., Ruttimann, U.E. and Unser, M. (1998) A pyramid approach to subpixel registration based on intensity. IEEE transactions on image processing : a publication of the IEEE Signal Processing Society 7: 27–41.

Triantaphylides, C., Krischke, M., Hoeberichts, F.A., Ksas, B., Gresser, G., Havaux, M., et al. (2008) Singlet oxygen is the major reactive oxygen species involved in photooxidative damage to plants. Plant Physiol 148: 960–968.

van Wijk, K.J. and Kessler, F. (2017) Plastoglobuli: Plastid Microcompartments with Integrated Functions in Metabolism, Plastid Developmental Transitions, and Environmental Adaptation. Annual Review of Plant Biology*, Vol* 68 68: 253–289.

Wada, S., Ishida, H., Izumi, M., Yoshimoto, K., Ohsumi, Y., Mae, T., et al. (2009) Autophagy plays a role in chloroplast degradation during senescence in individually darkened leaves. Plant Physiol 149: 885–893.

Wang, L.S., Leister, D., Guan, L., Zheng, Y., Schneider, K., Lehmann, M., et al. (2020) The Arabidopsis SAFEGUARD1 suppresses singlet oxygen-induced stress responses by protecting grana margins. Proc Natl Acad Sci U S A 117: 6918–6927.

Wang, S. and Blumwald, E. (2014) Stress-induced chloroplast degradation in Arabidopsis is regulated via a process independent of autophagy and senescence-associated vacuoles. Plant Cell 26: 4875–4888.

Wang, Y., Yu, B., Zhao, J., Guo, J., Li, Y., Han, S., et al. (2013) Autophagy contributes to leaf starch degradation. Plant Cell 25: 1383–1399.

Weaver, L.M. and Amasino, R.M. (2001) Senescence is induced in individually darkened Arabidopsis leaves, but inhibited in whole darkened plants. Plant Physiol 127: 876–886.

Woodson, J.D. (2019) Chloroplast stress signals: regulation of cellular degradation and chloroplast turnover. Curr Opin Plant Biol 52: 30–37.

Woodson, J.D., Joens, M.S., Sinson, A.B., Gilkerson, J., Salome, P.A., Weigel, D., et al. (2015) Ubiquitin facilitates a quality-control pathway that removes damaged chloroplasts. Science 350: 450–454.

Woodson, J.D., Perez-Ruiz, J.M. and Chory, J. (2011) Heme synthesis by plastid ferrochelatase I regulates nuclear gene expression in plants. Curr Biol 21: 897–903.

Ytterberg, A.J., Peltier, J.B. and van Wijk, K.J. (2006) Protein profiling of plastoglobules in chloroplasts and chromoplasts. A surprising site for differential accumulation of metabolic enzymes. Plant Physiol 140: 984–997.

Zhang, L., Kato, Y., Otters, S., Vothknecht, U.C. and Sakamoto, W. (2012) Essential role of VIPP1 in chloroplast envelope maintenance in Arabidopsis. Plant Cell 24: 3695–3707.

Zhuang, X. and Jiang, L. (2019) Chloroplast Degradation: Multiple Routes Into the Vacuole. Front Plant Sci 10: 359.

