## Supplementary material for "Singlet oxygen leads to structural changes to chloroplasts during degradation in the *Arabidopsis thaliana plastid ferrochelatase two* mutant": Fisher et al. 2021 supporting information

Figure S1: *fc2-1* mutants have increased singlet oxygen levels under 24h constant light conditions.

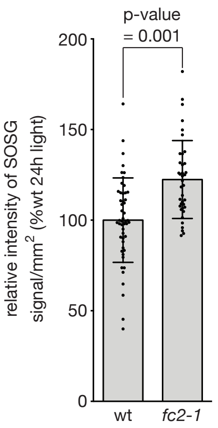


Quantification of relative singlet oxygen sensor green (SOSG) fluorescence in four-day-old seedlings grown under 24h constant light conditions. Shown are the mean SOSG fluorescence signal/mm$^2$ values +/ SD (n ≥ 38 seedlings). The statistical analysis was performed with a student's t test. Closed circles represent individual data points.

Figure S2: Visual phenotypes of seedlings.

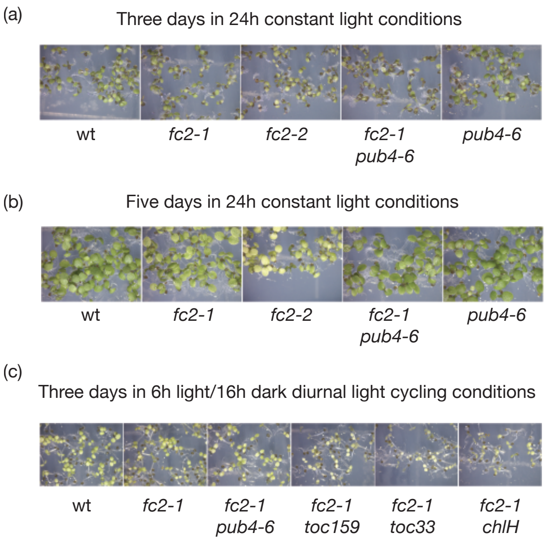


Shown are representative images of seedlings grown for **A)** three days under 24h constant light conditions, **B)** five days under 24h constant light conditions, and **C)** three days under 8h light/16h dark diurnal light cycling conditions. The genotypes of seedlings are indicated under the images.

Figure S3: A three dimensional analysis of chloroplast degradation within the central vacuole.

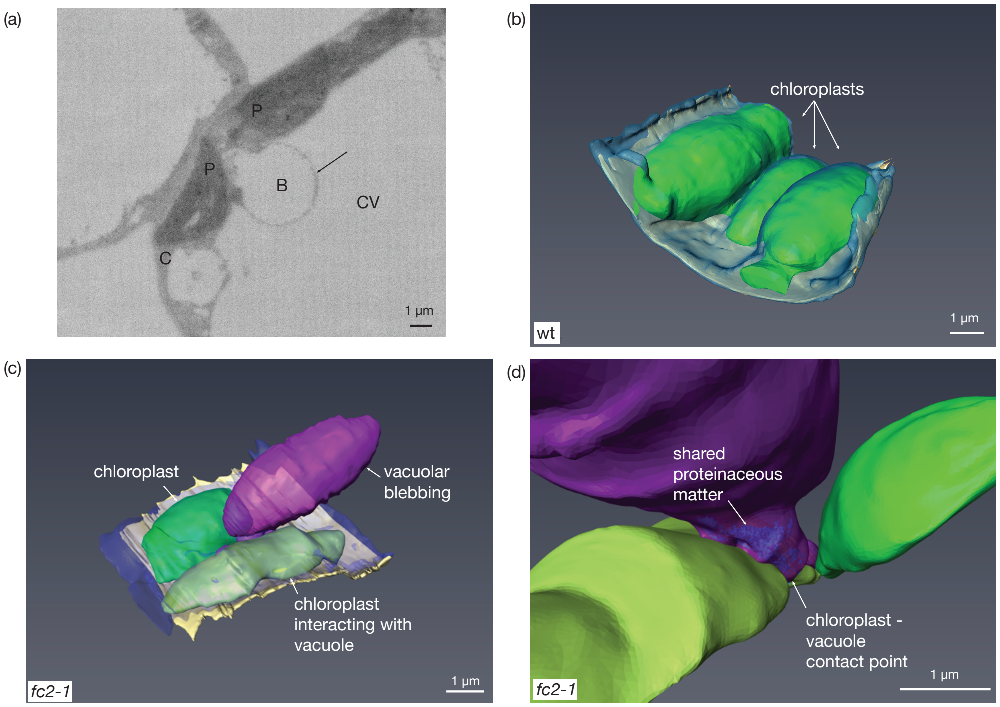


Chloroplast degradation in the *fc2-1* mutant was analyzed using three dimensional (3D) Serial Block Face-Scanning Electron Microscopy (SBF-SEM). **(A)** Shown is a snapshot of one unusually large vacuolar structure associating with a chloroplast (bleb volume, 177.6 $\mu m^3$; chloroplast volume, 87.9 $\mu m^3$, ratio of blub/chloroplast volume, 202%;, surface contact area, 7.4 $\mu m^2$). Scale bar = 1 µm. Abbreviations: CV, central vacuole, P, plastid; B, chloroplast/vacuole bleb; C, cytoplasm. Shown are 3D-renderings of **B**) three chloroplasts (in green) in wt, **C**) two chloroplasts in *fc2-1* from panel A, one of which is blebbing into the central vacuole (purple structure), and **D**) a zoomed-in image of the contact point between the vacuole and degrading chloroplast in panel C. Proteinaceous material shared between the two structures can be observed in dark blue.

<u>Figure S4: Assessment of cell death in *fc2-1* seedlings grown under different photoperiods.</u>

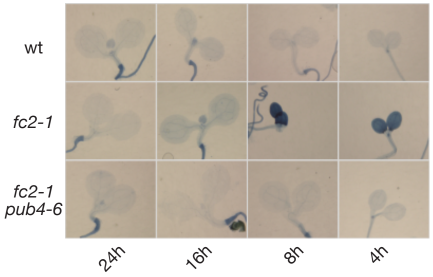


The effect of the photoperiod on chloroplast degradation was assessed. Shown are representative images of six-day-old seedlings grown under the indicated light conditions and stained with trypan blue. The dark blue color is indicative of cell death. Staining is quantified in Fig. 5a.

Figure S5: Assessment of plastoglobule association with thylakoid grana.

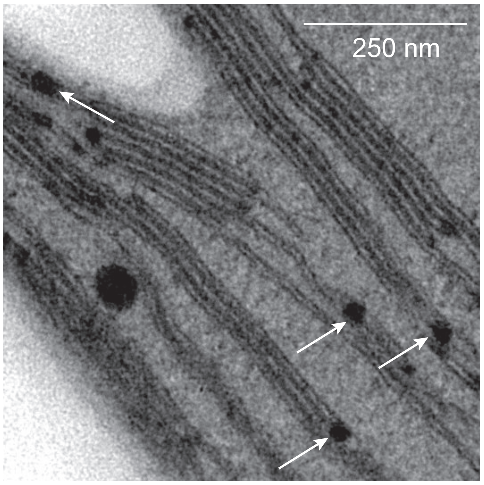


The association of plastoglobules with thylakoid grana membranes were assessed in various genotypes under different photoperiods. Shown are examples of plastoglubules associating with thylakoid grana in a five-day-old *fc2-1* seedling grown under 24h constant light. White arrows indicate plastoglobules that overlay with at least two thylakoid membranes. Scale bar = 250 nm.

**Table S1.**
Mutant plant lines used in study

| Mutant | Gene | mutation | notes | ref |
|---|---|---|---|---|
| *fc2-1* | *Plastid Ferrochelatase 2, FC2, At2g30390* | GABI_766H08 T-DNA in 5'UTR | Sulfadiazin[r] | (Woodson et al. 2011) |
| *fc2-2* | *Plastid Ferrochelatase 2, FC2, At2g30390* | SAIL_20_C06C T-DNA in intron7/9 | Basta[r] | (Woodson et al. 2011) |
| *pub4-6* | *Plant U-BOX 4, PUB4, At2g23140* | c9847535t, G255R | | (Woodson et al. 2015) |

**Table S2.**
Primers used in study

| Gene | Primer orientation / name | Sequence |
|---|---|---|
| **qPCR primer pairs** | | |
| *ACTIN2 At3g18780* | For. / JP199 | GCACTTGCACCAAGCAGCAT |
| | Rev. / JP200 | CCTTTCAGGTGGTGCAACGAC |
| *PPH, At5g13800* | For. / WLO1785 | ATGAGAAAGCCGGTTGTGAC |
| | Rev. / WLO1786 | AATGAACCAACGCCAAAGCC |
| *PES1, At1g54570* | Rev. / WLO1783 | AGCCAAGAAGAAGCAAAGCG |
| | For. / WLO1784 | ATGCCCATTGTCCTTGAAGC |

### References

Woodson, J.D., Joens, M.S., Sinson, A.B., Gilkerson, J., Salome, P.A., Weigel, D., et al. (2015) Ubiquitin facilitates a quality-control pathway that removes damaged chloroplasts. *Science* 350: 450-454.

Woodson, J.D., Perez-Ruiz, J.M. and Chory, J. (2011) Heme synthesis by plastid ferrochelatase I regulates nuclear gene expression in plants. *Curr Biol* 21: 897-903.
